# Convergent genomic signatures of high altitude adaptation among domestic mammals

**DOI:** 10.1101/743955

**Authors:** Dong-Dong Wu, Cui-Ping Yang, Ming-Shan Wang, Kun-Zhe Dong, Da-Wei Yan, Zi-Qian Hao, Song-Qing Fan, Shu-Zhou Chu, Qiu-Shuo Shen, Li-Ping Jiang, Yan Li, Lin Zeng, He-Qun Liu, Hai-Bing Xie, Yun-Fei Ma, Xiao-Yan Kong, Shu-Li Yang, Xin-Xing Dong, Ali Esmailizadeh Koshkoiyeh, David M Irwin, Xiao Xiao, Ming Li, Yang Dong, Wen Wang, Peng Shi, Hai-Peng Li, Yue-Hui Ma, Xiao Gou, Yong-Bin Chen, Ya-Ping Zhang

## Abstract

Abundant and diverse domestic mammals living on the Tibetan Plateau provide useful materials for investigating adaptive evolution and genetic convergence. Here, we utilized 327 genomes from horses, sheep, goats, cattle, pigs and dogs living at both high and low altitudes, including 73 genomes generated for this study, to disentangle the genetic mechanisms underlying local adaptation of domestic mammals. Although molecular convergence is comparatively rare at the DNA sequence level, we found convergent signature of positive selection at the gene level, particularly *EPAS1* gene in these Tibetan domestic mammals. We also reported a potential function in response to hypoxia for the gene *C10orf67*, which underwent positive selection in three of the domestic mammals. Our data provides insight into adaptive evolution of high-altitude domestic mammals, and should facilitate the search for additional novel genes involved in the hypoxia response pathway.

## Introduction

The high-altitude environments are considered physiologically challenging to endothermic animals. Plateaus, which occur on every continent and take up a third of the Earth’s land, are believed to provide fertile ground for investigating mechanisms of adaptation to hypoxia, and particularly convergent evolution (Storz et al. 2010; Cheviron and Brumfield 2012). To date, numerous studies have revealed special convergent physiological and morphological adaptations to hypoxia in many vertebrate species from highland. For example, endotherms living in high plateau environments have enhanced cardio-pulmonary function, increased sized hearts and lungs, as well as decreased morbidity due to pulmonary diseases compared with closely related species from lowland environments (Qiu et al. 2012). Tibetans were found to exhibit characteristics contributing to improved hypoxic and hypercapnic ventilatory responsiveness including larger lung volumes, better pulmonary function, and greater lung diffusing capacity (Wu and Kayser 2006). They also maintain higher levels of arterial oxygen saturation both at rest and during exercise, and show reduced loss of aerobic performance with increasing altitude (Wu and Kayser 2006). Several aspects of the genetic basis for the adaptation to hypoxia have been elucidated in some vertebrate species (Storz et al. 2007; Storz et al. 2009; Beall et al. 2010; Simonson et al. 2010; Yi et al. 2010; Wilson and Hay 2011; Qiu et al. 2012; Ge et al. 2013; Li et al. 2013; Qu et al. 2013; Foll et al. 2014; Gou et al. 2014; Li et al. 2014; Lorenzo et al. 2014; Wang et al. 2014; Zhang et al. 2014; Natarajan et al. 2015; Tufts et al. 2015). In particular, these studies suggest that modifications of hemoglobin function is key to mediating an adaptive response to high-altitude hypoxia in a number of mammals, birds, and amphibians (Storz 2007; Storz and Moriyama 2008; Storz et al. 2009; Natarajan et al. 2015; Tufts et al. 2015). Despite this, much remains to be learned about the molecular genetic basis underlying high-altitude adaptation in general. To achieve this goal, analyzing whether molecular convergent evolution occurs commonly for high altitude adaptation among different species may provide important insights.

The Tibetan Plateau, with an average elevation of over 4500 m, is the largest and highest plateau in the world. The extremely harsh environment of this plateau includes low oxygen content, low air pressure, low temperatures and strong UV radiation. Indeed, the unique geographical and environmental features have shaped the biological diversity of the Tibetan Plateau, and provide fertile ground for investigating the genetic loci underlying adaptation to high altitude in humans, in wild animals such as chiru and Tibetan wolf, as well as in domestic animals like yak, Tibetan chicken, Tibetan Mastiff (dog) and Tibetan pig (Beall et al. 2010; Simonson et al. 2010; Yi et al. 2010; Wilson and Hay 2011; Qiu et al. 2012; Ge et al. 2013; Li et al. 2013; Qu et al. 2013; Foll et al. 2014; Gou et al. 2014; Li et al. 2014; Lorenzo et al. 2014; Wang et al. 2014; Zhang et al. 2014). Previous findings have shown that animals primarily domesticated at low altitudes (sheep, goats, cattle, chickens, donkeys, pigs, dogs and horses) have been brought to the Tibetan plateau in the past few thousand years by Tibetans, which likely facilitated the permanent human occupation of this plateau (Chen et al. 2015). Although these Tibetan domestic animals have inhabited the plateau for less than 4000 years (Chen et al. 2015), they harbor special physiological traits that allow their survival in Tibet. Studying the genetic basis of these high-altitude adaptation in these animals will not only promote our understanding of their inhabitation history, but also provide essential information about the molecular convergent evolution process.

In the present study, to reveal the genetic mechanisms underlying high-altitude adaptation of domestic mammals, we utilized genomes, with an additional 73 generated for this study, from horse, sheep, goat, cattle, pig and dog populations that live in the Tibetan Plateau and at low altitudes. We investigated positive selection operating on these Tibetan domestic mammals to detect the occurrence of convergent evolution for high-altitude adaptation.

## Results

In the present study, a total of 327 (134 individuals inhabiting the Tibetan Plateau including 19 goats, 24 horses, 20 sheep, 11 dogs, 19 cattle and 41 pigs, and 193 control individuals from lowland areas including 20 goats, 10 horses, 70 sheep, 34 dogs, 30 cattle and 29 pigs, **Supplementary Table S1**) whole genome sequences (including 73 new sequences) were analyzed to identify genome-wide SNPs. To explore potential adaptation to high altitude, differentiation at SNPs in each species was calculated using XP-EHH (cross population extended haplotype homozygosity) (Sabeti et al. 2007), ΔDAF (the difference of the derived allele frequencies), and *F*_ST_ (Wright 1965; Akey et al. 2002) between the highland and lowland populations to evaluate their evolutionary properties.

### Patterns of convergent evolution at the DNA sequence level

We first studied potential convergent evolution at the DNA sequence level. SNPs from each domestic mammal were coordinated to the human genome (hg19) using the Liftover program (https://genome.ucsc.edu/cgi-bin/hgLiftOver). Through this analysis, we found an average of 182K SNPs occurring at the same human coordinated position for any pair of domestic mammals (**Supplementary Table S2**). We retrieved SNPs showing values of *F*_ST_, XP-EHH and ΔDAF higher than the 99th percentile values of genome wide SNPs, and found the proportion of SNPs showing higher values is nearly 1% among these SNPs occurring at the same human coordinated position for any pair of domestic mammals (**Supplementary Table S3**). We further identified potential convergent site, at which SNPs showing higher values of *F*_ST_, XP-EHH or ΔDAF in both species for any pair comparison. This analysis identified only 3-45 sites showing potential convergent evolution between any pair of Tibetan domestic mammals (**Supplementary Table S4**). This analysis supports the previous finding that molecular convergence at the DNA sequence level is not frequent (Bigham et al. 2015; Foote et al. 2015; Zou and Zhang 2015).

### Shared signatures of positive selection in Tibetan domestic mammals at the gene level

We integrated the *F*_ST_, XP-EHH and ΔDAF analyses (named *iFXD*) as previously described (Grossman et al. 2013) to improve our power and to identify positively selected genes in the six Tibetan domestic mammals. Based on the outlier approach in the top 1% ranking, in total, 199, 187, 190, 181, 189 and 170 protein coding genes were identified to exhibit extreme divergence with evidence for positive selection in goats, horses, sheep, dogs, cattle and pigs, respectively (**Supplementary Table S5**). After inference of demographic history and population simulation, we found that the outlier approach is robust to demographic history and SNPs density (**Supplementary Note, Figs. S1-S4, Table S6-S10**). Among these, 43 protein coding genes were found to display the evidence for positive selection in at least two lineages (Fig. 1A, **Supplementary Fig. S5**). To demonstrate that these were not false-positives, we conducted a permutation test by randomly sampling genes. In all 1000 random sampling, the numbers of shared positively selected genes was less than 43 (Fig. 1A). We further investigated evidence of shared positive selection in a set of one-to-one orthologous genes by integrating signals of selection among the six Tibetan domestic mammals. We retrieved 30 genes that had potentially evolved under positive selection (in the top 5% ranking) in at least three mammals (**Supplementary Fig. S6**). When we examined 1000 sets of randomly chosen genes, none of the datasets contained more than 20 genes under positive selection in at least three mammals (Fig. 1B). These data indicate that convergent signature of positive selection occurred among these different species.

**Figure 1:**
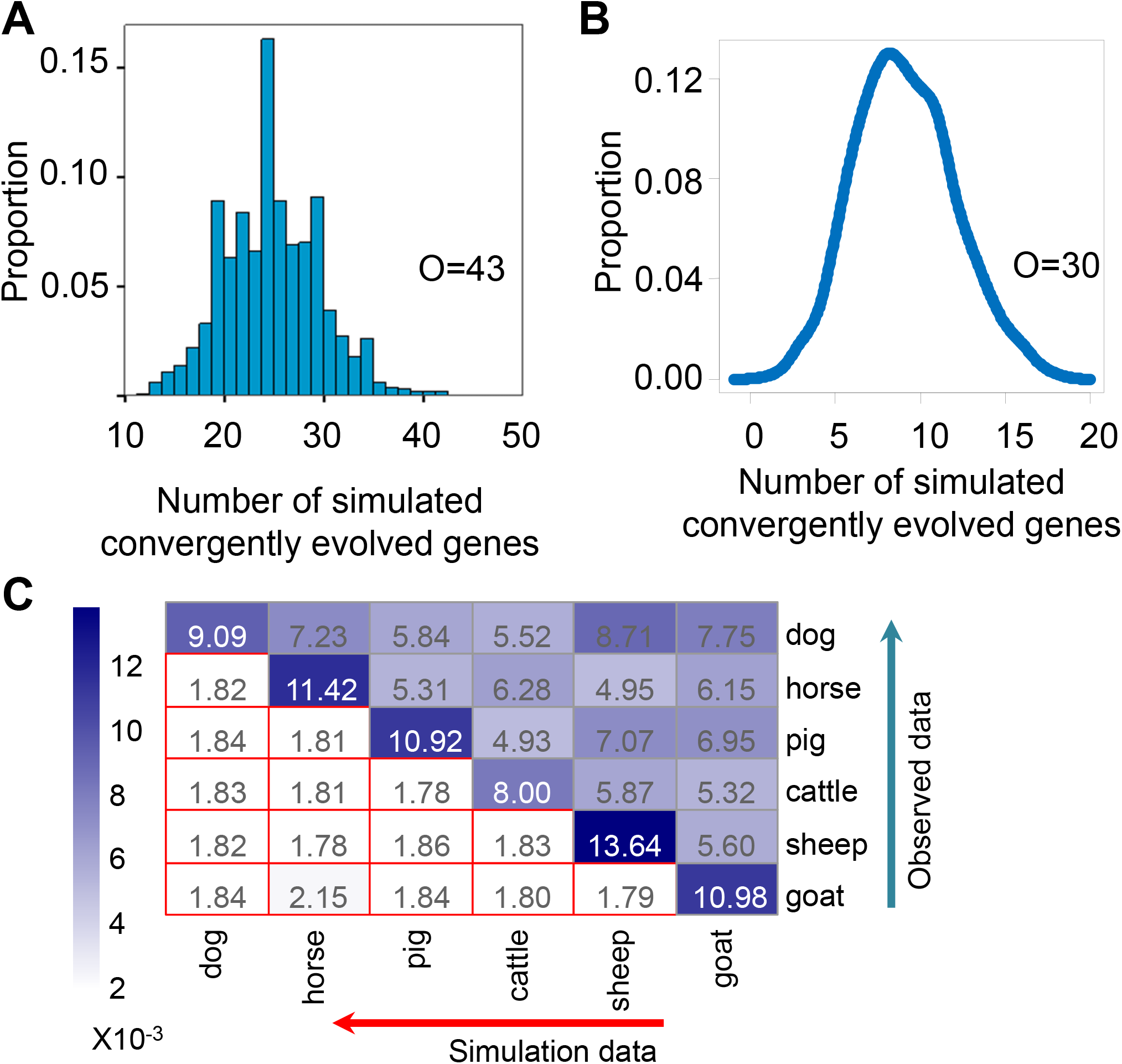
Frequent shared targets of positive selection in Tibetan domestic mammals. (A) Distribution of the numbers of simulated shared positively selected genes. Observed number of shared positively selected genes was 43 (O=43). Namely, 43 protein coding genes evolved under positive selection in at least two Tibetan domestic mammals (in the top 1% ranking). (B) Distribution of the numbers of simulated shared positively selected genes based on one-to-one orthologous genes. We further investigated evidence of convergent signature of positive selection in 10103 one-to-one orthologous genes. The number of genes observed to potentially evolve under positive selection (in the top 5% ranking) in at least three mammals is 30 (O=30) (Supplementary Fig. S2). (C) Observed and simulated ratios of gene-gene interactions among positively selected genes within species and between positively selected genes of different species. Lower red circled boxes present the simulated ratios, and above blue boxes are observed ratios (×10^−3^). Detail of the simulations are provided in the SI Appendix.

We then took one further step to examine whether positive selection was occurring on particular single genes or on genes within certain pathways. For this, we conducted a gene enrichment analysis, and found that a gene-gene interaction network containing *EPAS1* was overrepresented in the candidate positively selected genes in each of the Tibetan mammals examined (**Supplementary Fig. S7**). *EPAS1* encodes a subunit of the HIF transcription factor and is a key gene for hypoxia adaptation in Tibetans (Beall et al. 2010; Simonson et al. 2010; Yi et al. 2010). In our analyses, this gene was found to have evolved under positive selection in five Tibetan mammals (top 1% ranking) and in Tibetan cattle (top 5% ranking), corroborating the previously reported convergence of genetic adaptation to high altitude in dogs and humans (Beall et al. 2010; Simonson et al. 2010; Yi et al. 2010; Gou et al. 2014; Li et al. 2014; Wang et al. 2014). These data suggested that during adaptive evolution, positive selection drove changes in genes within a network involved in similar functional pathways rather than single genes.

Inspired by the above findings, we further investigated whether positively selected genes exhibited more frequent interactions with each other among different species. As expected, positively selected genes displayed significantly more gene-gene interactions with each other than randomly selected genes (Fig. 1C). In addition, positively selected genes of one species also interacted significantly with those of other species based on the gene-gene interaction data from human (Fig. 1C).

### Common rapid evolution of hypoxia response genes in Tibetan domestic mammals

To validate the contention that positive selection drove changes in genes within a network involved in similar functional pathways, we examined the evolution of genes involved in the hypoxia response pathway, which might play an important role in high altitude adaptation. The hypoxia response genes were defined in the study (Huerta-Sanchez et al. 2013), which used the AmiGO tool (http://amigo.geneontology.org) retrieve all genes within the Gene Ontology biological process term “response to hypoxia” plus all descendent terms (GO:0001666 “response to hypoxiam” GO:0071456 “cellular response to hypoxia” and GO:0070483 “detection of hypoxia”). *F*_ST_, XP-EHH and ΔDAF analyses got different results, which is likely due to different statistical principles these methods used, as described in the previous study (Sabeti et al. 2006). However, as seen in the values of *F*_ST_, XP-EHH and ΔDAF, SNPs in a set of hypoxia response genes displayed significantly higher levels of differentiation compared with SNPs in the remaining genome-wide set of genes (*P*<0.01, by Mann-Whitney U test, Fig. 2). To further validate the significance and exclude the confounding factor of different numbers of genes for comparison, we randomly selected a group of genes with the same number with hypoxia response genes for each species. As expected, we got similar significant results (*P*<0.01, Fig. 2). In addition, SNPs within genes in the gene ontology (GO) category associated with hypoxia also harbored significantly higher values for *F*_ST_, XP-EHH and ΔDAF (**Supplementary Fig. S8**). These lines of evidence indicated common rapid evolution of hypoxia response genes, allowing adaptation to hypoxia in these Tibetan domestic mammals.

**Figure 2:**
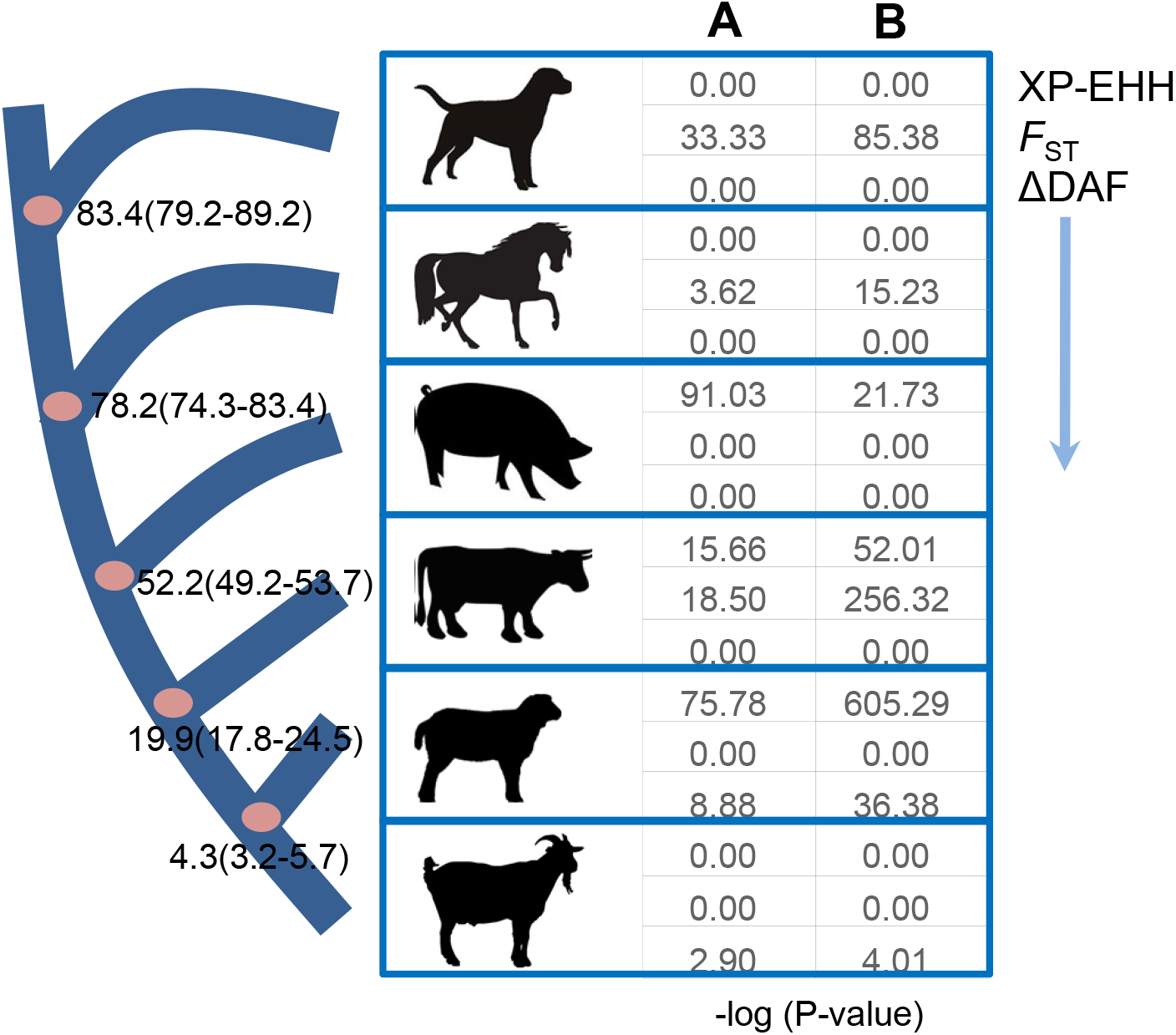
Rapid evolution of hypoxia response genes in six Tibetan domestic mammals. Significantly higher values of XP-EHH (cross population extended haplotype homozygosity), *F*_ST_ and ΔDAF (the difference of the derived allele frequencies) were observed for SNPs found in genes known to be involved in hypoxia response than in other genome-wide genes or randomly selected genes. Column A (compared with genome-wide genes) and B (compared with random genes) represent the –log10 transformed *P*-values calculated using Mann–Whitney U tests. The hypoxia response genes were retrieved from a previous study (Huerta-Sanchez et al. 2013). Numbers on the nodes of the phylogenetic tree are the divergence times from present (millions of years) and their confidence intervals. Dates are from a previous study (Jiang et al. 2014).

### Convergent signature of positive selection on *EPAS1* for attenuating hemoglobin concentration in Tibetan humans and Tibetan horses

The above finding suggested that, although signatures of positive selection were significantly shared among different species, the sites targeted by selection in genes under convergent evolution are not identical. We reasoned that positive selection on the same genes might generate convergent functions or phenotypes in different species even though the sites targeted by selection were different. To validate this, we examined *EPAS1*, which is under positive selection in Tibetan humans and Tibetan domestic mammals, to determine whether changes at different sites in different species generated similar functional consequences. In Tibetan humans, adaptive variants of *EPAS1* favors adaptation to high altitude by attenuating maladaptive physiological alterations such as elevated hemoglobin concentration (Bigham and Lee 2014). We choose a SNP (chr15: 52623471), which harbors the highest *iFXD* value in horse *EPAS1* and tested its association with hematologic traits in the Tibetan horse. As expected, we saw that the *EPAS1* SNP showed significant correlations with some measured physiological traits (**Supplementary Fig. S9**, P<0.01 after false discovery rate (FDR) correction). Consistent with Tibetan humans, the derived allele of *EPAS1* SNP (chr15: 52623471) in the Tibetan horse was associated with lower hemoglobin concentration (HGB) (**Supplementary Fig. S9**), and its frequency in the Tibetan and lowland horses was 0.79 and 0.28, respectively. One may argue that the association may be attributable to demographic history, like frequent introgression from lowland horse. Under demographic history, SNPs across the genome showing differentiation between high and low populations would correlate with these blood phenotypes. Therefore, we further chose a SNP which changes the amino acid of EPAS1 (R144C), and harbors the largest *iFXD* value among all of the non-synonymous SNPs examined. However, it didn’t show association with any of the blood-related traits. We proposed that the correlation of *EPAS1* SNP (chr15: 52623471) with blood traits is likely attributable to adaptive evolution, although we can’t absolutely exclude the possibility of demographic history.

Increased hemoglobin concentration and erythrocytosis after exposure to high altitude are counterproductive for individuals from lowlands, as these changes result in increased blood viscosity that reduces uterine artery blood flow and a reduced rate of O_2_ delivery to the uteroplacental circulation (Storz et al. 2010), which ultimately lead to elevated risk of stillbirth, preterm birth and reduced birth weight. In addition, individuals from low altitudes exposed to high altitudes often develop high hemoglobin concentrations associated with disadvantageous physiological outcomes including hemodynamic hyperviscosity, cardiac dysfunction and chronic mountain sickness (CMS) (Bigham and Lee 2014). Convergent evolution of *EPAS1* potentially occurred in Tibetan domestic mammals and Tibetan humans to attenuate maladaptive physiological alterations such as elevated hemoglobin concentration and erythrocytosis.

### Function of a positively selected gene *C10orf67* in the response to hypoxia

Since genes known to be involved in the response to hypoxia had evolved under positive selection in Tibetan domestic mammals, we reasoned that novel genes involved in the hypoxia response pathway might be identified from our list of positively selected gene candidates. To test this, we examined the role of *C10orf67*, a gene with uncertain function but displayed evidence of positive selection in three Tibetan mammals (**Supplementary Figs. 5, 6**), in cellular response to hypoxia. Hypoxia will induce apoptosis, a form of programmed cell death. It is necessary to detect the apoptosis under hypoxia condition to reveal the role of *C10orf67* in the response to hypoxia. We examined the rate of apoptosis in NIH-3T3 cells stably expressing two independent shRNAs targeting *C10orf67* under normoxic (21% O_2_) and hypoxic (1% O_2_) conditions (Fig. 3A-3B). In contrast to the increased rate of apoptosis observed in control cells transfected with scrambled shRNA under hypoxia compared with normoxia, *C10orf67*-deficient cells displayed a dramatic reduction in the rate of apoptosis under hypoxic condition (Fig. 3B-3C). We then used whole transcriptome sequencing (RNA-seq) to characterize the regulatory network involving *C10orf67*. We compared the results obtained from cells depleted of *C10orf67* to those from cells transfected with scrambled shRNA under both normoxic and hypoxic conditions and observed 263 and 206 differentially expressed (differential expression was defined by having at least a 2-fold change) genes, respectively (*P*<0.01, Fig. 3D). Intriguingly, many of the identified genes are involved in the response to changes in oxygen levels, angiogenesis, cell death and apoptotic processes (Fig. 3E, **Supplementary Table S11**), indicating a potential role for *C10orf67* in the response to hypoxia or during tumorigenesis.

**Figure 3:**
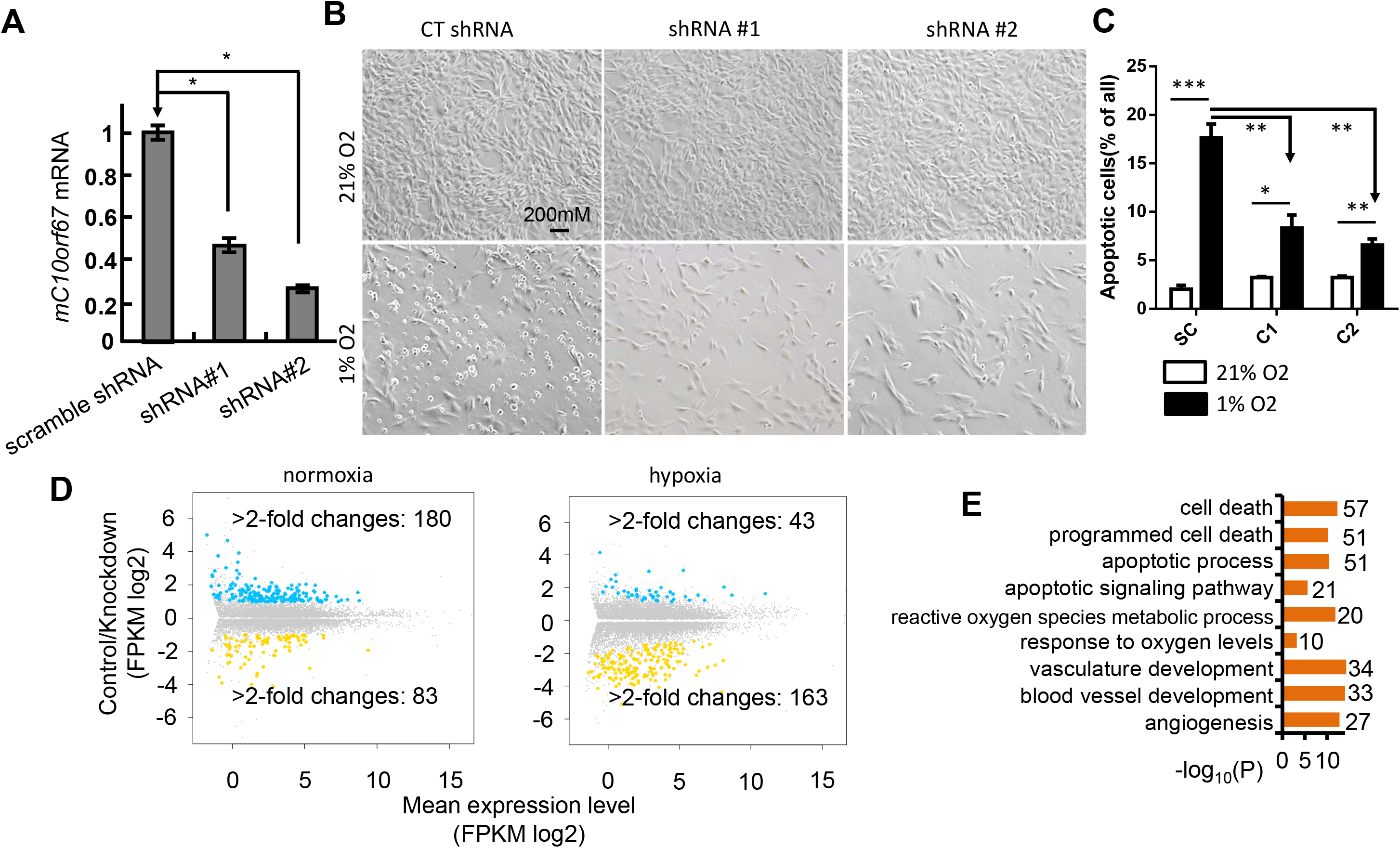
Functional analysis of *C10orf67*. (A) Relative mRNA expression levels of *mC10orf67* in NIH-3T3 cells stably expressing two independent shRNAs compared with scramble-control shRNA. \**P*<0.05. (B) Suppression of apoptosis by depleting *C10orf67* expression in NIH-3T3 cells under 1% O_2_ hypoxic conditions. (C) After 72 h treatment under hypoxic conditions, NIH-3T3 cells were stained with annexin V/PI, and the percentage of apoptotic cells was assessed by flow cytometry. The scramble-control shRNA stable cell line served as a control. SC, C1 and C2 indicate cell treated by control shRNA, shRNA #1, and shRNA#2 respectively. \**P*<0.05, \*\**P*<0.001. (D) Comparison of gene expression with scramble-control and *C10orf67* knockdown by RNA-seq under normoxic (left panel) and hypoxic (right panel) conditions. A summary dot plot is shown, with blue dots representing down-regulated and yellow dots representing up-regulated genes (*n*=3 samples for each, at least 2-fold difference and *P*-corrected < 0.05). (E) List of GO (gene ontology) terms related to hypoxia and apoptosis that were significantly enriched among the differentially expressed genes between the scramble-control and *C10orf67* knockdown cells under normoxic conditions by RNA-seq. The *x*-axis shows the –log10 transformed FDR corrected *P*-values. The number of enriched genes in each GO is also presented.

### Positive selection drove the population differentiation of non-synonymous SNPs in Tibetan sheep, dog and cattle

Local positive selection tends to increase population differentiation, and preferentially targets functional SNPs resulting in amino acid change or cis-regulatory events, while under the assumption of neutrality, population differentiation is due to demographic history, all loci are affected similarly (Barreiro et al. 2008). In the current study, population differentiation, which was evaluated by *F*_ST_ (Akey et al. 2002), was more pronounced at non-synonymous SNPs than any other types of SNPs (e.g. SNPs located at the upstream/downstream of genes, intronic SNPs, synonymous SNPs) in sheep, dogs, cattle and horses (**Supplementary Fig. S10-S11, Supplementary Table S12**). Although local adaptation tends to increase population differentiation, many studies have also argued that background selection reduces the within-population diversity, and should lead to high *F*_ST_ (Cutter and Payseur 2013; Cruickshank and Hahn 2014; Hoban et al. 2016). We further calculate the derived allele frequencies of these high *F*_ST_ non-synonymous SNPs, and found that they harbor significantly higher derived allele frequencies in Tibetan domestic mammals, like sheep, dog and cattle (**Supplementary Fig. S12**). Therefore, positive selection likely has driven population differentiation of functional SNPs between Tibetan sheep, dog and cattle and other lowland populations, and likely contributes to local adaptation. For example, highly differentiated non-synonymous SNPs occurred in *EPAS1* of Tibetan dogs and cattle (**Supplementary Table S13**).

We further examined non-synonymous SNPs harboring high levels of population differentiation between Tibetan domestic mammals and lowland mammals (*F*_ST_>0.5) within these positively selected genes in each species (**Supplementary Table S13**). We found that these genes were mainly involved in apoptosis, metabolism and functions of the nervous system, supporting the widely accepted tight connection between hypoxia adaptation and these pathways (Greijer and van der Wall 2004; LaManna 2007; Masson and Ratcliffe 2014; Eales et al. 2016). However, the functional consequences of these amino acid changing mutations remain to be determined. Functional validation of the association between these mutations and high-altitude adaptation will provide essential information about potential biomarkers for breeding at high altitude.

### Other genes under positive selection in different domestic mammals

In the Tibetan sheep, we found that *MITF* (melanogenesis associated transcription factor) evolved under positive selection. *MITF* is crucial for pigmentation by regulating the differentiation and development of melanocytes (Levy et al. 2006). The factor driving positive selection on *MITF* and whether it is related to high altitude adaptation is unclear, but is likely to be associated with phenotypic evolution in Tibetan sheep. Indeed, while Tibetan sheep often have a white coat, 90% of the total population have variegated wool covering their heads and limbs, and solid-white or solid-black sheep are rare (Han et al. 2005). The positive selection observed for *MITF* is therefore likely associated with the change in coat pigmentation found in the Tibetan sheep. This variegated wool covering the heads and particularly eyes of the Tibetan sheep is believed to help them resist the strong UV radiation in highland areas (Han et al. 2005).

In the Tibetan goat, the gene *DSG3* harbors the most significant signal of positive selection. A sliding window analysis (window size: 50kb, step size: 25kb) of the *iFXD* values across the whole genome also revealed a region with significant values that spanned the *DSG* gene cluster (including *DSG2*, *DSG3*, and *DSG4* genes) (**Supplementary Fig. 13**). *DSG* encode desmosome proteins that are involved in the intercellular junctions of epithelia and cardiac muscle cells, and play important roles in the development of the heart, as shown by mutations within desmosome that cause Arrhythmogenic right ventricular cardiomyopathy in humans (Delmar and McKenna 2010). Endothermic animals living on high plateaus typically have strong cardio-pulmonary function with larger hearts. The important role of *DSG* genes in heart development raised the possibility that positive selection on these genes might have yielded phenotypic changes in the Tibetan goat that were helpful for high altitude adaptation. However, the caveat is that the underlying factor driving positive selection on gene *DSG* is not only high altitude adaptation. For example, genes *DSG* also have important function in immune system (Amagai et al. 1998), which is also the driving factor for rapid evolution of genes.

RYR2 (ryanodine receptor 2), functions as a component of calcium channels that supplies Ca^2+^ for cardiac muscle excitation-contraction coupling. Previous studies have reported evidence of positive selection on *RYR2* in the Tibetan chicken (Wang et al. 2015) and Tibetan wolf (Zhang et al. 2014), presumably for a role in adaptation to hypoxia. Here, we also found evidence of positive selection on *RYR2* in the Tibetan goat. Hypoxia should induce the release of reactive oxygen species (ROS) and cause an influx of extracellular Ca^2+^ mediated by ion channels as well as the intracellular release of Ca^2+^ from stores by the ryanodine receptor. Increased intracellular Ca^2+^ will then cause contraction of cells, an important response to hypoxia (Hui et al. 2006; Shimoda and Undem 2010; Wang and Zheng 2010).

In Tibetan cattle, we observed positive selection on the gene *HMGA2*. Variants within *HMGA2* have been reported to be associated with body height in humans and domestic animals (Weedon et al. 2007; Weedon et al. 2008; Makvandi-Nejad et al. 2012; Kader et al. 2015). In our results, a nonsynonymous SNP (A64P) within *HMGA2* displayed high level of population differentiation between Tibetan cattle and other cattle (**Supplementary Table S13**). In addition, *ADH7* also evolved under positive selection in the Tibetan cattle, and harbored three non-synonymous SNPs showing high level of differentiation between Tibetan cattle and other cattle. *ADH7* encodes a class IV alcohol dehydrogenase that is very active as a retinol dehydrogenase. The gene had been reported to evolve under positive natural selection in Tibetan humans, and was associated with their weight and body mass index (BMI) (Yang et al. 2017). In sum, we believe that positive selection on *ADH7* might be a potential convergent signature, and be associated with the stature of Tibetan cattle.

Several recent whole genome sequencing studies have examined high-altitude adaptation of Tibetan pigs (Li et al. 2013; Ai et al. 2014; Dong et al. 2014). To further explore potential genetic events for adaptation of Tibetan pigs, we performed RNA-sequencing to profile the transcriptomes of lung tissues from four Tibetan pigs and four Min pigs collected at a low altitude. Four genes *ADAMTSL4*, *ZFAND2A*, *IER2*, and *OVOS2* were found to evolve under positive selection and exhibited significant down-regulated mRNA expression in the Tibetan pigs (**Supplementary Fig. S14**). Whether down-regulation of these genes contributes to high altitude adaptation of Tibetan pigs still remains unclear, however, the fact that *ADAMTSL4*, *IER2* and *OVOS2* might participate in the process of angiogenesis (Le Goff and Cormier-Daire 2011; Huang et al. 2013; Wu et al. 2015) supports their crucial roles in hypoxia adaptation.

## Discussion

### Shared targets of positive selection for local adaptation of domestic mammals

In the present study, we utilized large-scale genomic data to acquire insights into the adaptive evolution of Tibetan domestic mammals. Our results were indicative that targets of positive selection were commonly shared among different species. It may suggest convergent evolution occurring among different domestic mammals. However, we didn’t find frequent molecular convergence at the DNA level. Therefore, convergent signature of positive selection could also induce the rapid divergence of genes among these species.

One caveat of study on convergent evolution is false positive (Bigham et al. 2015; Foote et al. 2015; Zou and Zhang 2015; Xu et al. 2017). Signal of convergent evolution may be observed frequently between control groups, such as lowland populations in this study. Therefore, more populations from lowland will be needed in the future to identify positively selected genes in the lowland populations and be helpful for studying pattern of convergent evolution in the control population. Another caveat of our study is that the factors driving positive evolution are unclear. The parsimony factor is high altitude adaptation. However, we also note that some positively selected genes are involved in immune system, which have been found to evolve rapidly in many organisms due to arm race hypothesis. Therefore, common rapid evolution of immune genes might also be a factor for shared signature of positive selection.

In addition, we also report the likely role of a positively selected gene *C10orf67* in the response to hypoxia. We reasoned that studying genetic loci underlying high-altitude adaptation should facilitate the identification of novel genes in unique pathways such as hypoxic response.

### Why is positive selection on *EPAS1* a common event in all these mammals?

Utilizing large-scale genomic data, we reported the putative contribution of multiple positively selected genes, particularly *EPAS1*, and positive selection on amino acid changing SNPs, to high-altitude adaptation of Tibetan domestic mammals. While we have acquired insights into the adaptive evolution of these mammals, which variants at *EPAS1* facilitate high-altitude adaptation of Tibetan domestic mammals remains unclear. Therefore, more experiments dissecting the functional consequences of these variants are needed.

It is assumed that phenotypic traits shared by Tibetan domestic mammals are likely attributed to high-altitude adaptation changes associated with positive selection on certain genes, and *EPAS1* is likely one of those. Elevated hemoglobin concentration and erythrocytosis occur commonly during the initial short-term acclimatization responses to hypoxia. While these physiological changes affect their breeding and health (Storz et al. 2010; Bigham and Lee 2014), those eventually adapted must have undergone a series of physiological processes to maintain homeostasis. Therefore, one of the reasons for frequent positive selection on *EPAS1* might be to attenuate the rapidly induced common elevated hemoglobin concentration and erythrocytosis (Storz et al. 2010; Bigham and Lee 2014). Given the fact that *EPAS1* encodes one of the subunits of the hypoxia-inducible factors (HIFs) and has pleiotropic functions, it is of great interest to determine the functional consequences of selection on *EPAS1* and associated physiological processes in future studies.

### Potential application of adaptive mutations in domestic mammal breeding at high altitude

The extreme environment of the Tibetan plateau limits livestock husbandry, which dominates the Tibetan economy. To solve the problem that many of the domestic animals transported from lowland areas cannot live or reproduce well at high altitude, we sought to identify variants in genes underlying hypoxia adaptation in domestic animals so that commercial breeds of domestic mammals from lowland areas carrying advantageous variants can be generated and introduced to different plateaus. It is intriguing to confirm that *EPAS1* evolved under positive selection for hypoxia adaptation, and we believed that certain functional variants within *EPAS1* could be used to help improve livestock breeding. In addition, many mutations related to amino acid changes showed great population differentiation between highlanders and lowlanders, suggesting their potential as the candidate biomarkers for economic purposes, although further studies are needed to validate their biological roles in high-altitude adaptation.

## Materials and Methods

### Genome sequences and SNPs calling

In this study, blood samples from 19 goats, 20 sheep, and 34 horses were collected from individuals inhabiting different regions of the Tibetan Plateau and surrounding lowland regions in China. DNA was extracted from the blood samples using a standard Phenol–chloroform extraction protocol. Using the Illumina standard protocol, genomic libraries with insert size ~400 bp were constructed, and sequenced by Hiseq2000 or Hiseq2500. Additional genomes were from other studies (Gou et al. 2014; (!!! INVALID CITATION !!!). A final total of 134 genomes from Tibetan domestic mammals (19 Tibetan goats, 24 Tibetan horses, 20 Tibetan sheep, 11 Tibetan Mastiffs, 19 Tibetan cattle and 41 Tibetan pigs) and 193 genomes of domestic mammals from lowlands were used for population genomic analysis (Supplementary Table S1).

Reference genome sequences of cattle, horse, pig, dog and sheep were downloaded from ENSEMBL (version 72), and the reference genome sequence of the goat was download from http://goat.kiz.ac.cn/ (CGD) (Dong et al. 2013). Low-quality bases in sequence reads were filtered using Btrim, and qualified reads were aligned onto the references using BWA-MEM with default settings except for “-t 8 -M -R” options (Li 2013). A series of post-processes were carried out with the aligned bam format files, including sorting, duplicates marking and local realignment using the relevant tools in the Picards (picard-tools-1.56) and Genome Analysis Toolkit (GATK, GenomeAnalysisTK-2.6-4-g3e5ff60) packages (McKenna et al. 2010). SNPs were called and filtered using the UnifiedGenotyper and VariantFiltration commands in GATK. Loci with RMS mapping quality less than 25 and genotype quality less than 40, for which reads with zero mapping quality constituted more than 10% of all reads at this site, were removed. Loci with more than two alleles and within clusters (more than three SNPs in a 10 base pair (bp) window) were removed. All SNPs were annotated using the ANNOVAR program (www.openbioinformatics.org/annovar/), which annotated different types including non-synonymous, synonymous, splicing, 5’ UTR, 3’ UTR, intronic, upstream, downstream and intergenic SNPs. Variants that were missing with <50% were imputed and haplotypes were inferred using BEAGLE (http://faculty.washington.edu/browning/beagle/beagle.html) (Browning and Browning 2007).

### Genes associated with hypoxia

Sequences of 154 human genes involved in the hypoxia response pathway were downloaded from a previous study (Huerta-Sanchez et al. 2013). One-to-one orthologous genes between humans and domestic mammals, including dog, pig, horse, cattle and sheep, were downloaded from Ensembl using BioMart (http://www.ensembl.org/). One-to-one orthologous genes between humans and goat were retrieved by a reciprocal best-hit BLASTP search.

### Calculation of *F*_ST_, XP-EHH, ΔDAF, and *iFXD*

We used *F*_ST_ (Wright 1965; Akey et al. 2002), XP-EHH (Sabeti et al. 2007) and ΔDAF to evaluate the differentiation of each SNP between the highland and lowland populations in each mammal. The *F*_ST_ of each SNP was calculated as previously described (Akey et al. 2002). *F*_ST_ with negative values having no biological explanation and were arbitrarily set to 0. The XPEHH program (http://hgdp.uchicago.edu/Software/) was used to calculate the XP-EHH value for each SNP. Here, we did not consider variation of the recombination rate due to the lack of data for these genomes. However, the EHH method, which does not consider variation in the recombination rate, has been proven to possess the power to detect signatures of positive selection, and XP-EHH has been successfully used to detect positively selected genes in several species including dogs (Li et al. 2014), chickens (Wang et al. 2015) and horses (Kader et al. 2015) that lack recombination rate data. ΔDAF was calculated for each SNP as the DAF in the domestic animals from the highlands minus the DAF in domestic animals from the lowlands. The ancestral allele was inferred according to the allele of the outgroup, including the European pig for Tibetan pig, yak for Tibetan cattle, golden jackal for Tibetan dog, Przewalski’s horse for Tibetan horse, sheep for Tibetan goat and *Ovis dalli* for Tibetan sheep.

We calculated *iFXD* for each SNP by integrating *F*_ST_, XP-EHH and ΔDAF. 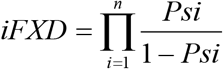, where *i* is the three different methods and *P*s represents the probability that the SNP is under positive selection.

### Identification of candidate genes under positive selection in the Tibetan domestic mammals based on *iFXD* values

The *iFXD* value of each protein coding gene was calculated by averaging the *iFXD* values of all SNPs within a gene. The demographic history (a genome-wide force that affects patterns of variation at all loci in a genome in a similar manner, whereas directional selection acts upon specific loci (Sabeti et al. 2006)) of domestic mammals in Tibet might be much more complex due to genetic admixture between mammals from highland and lowland areas. Inferring and simulating the demographic history of these Tibetan domestic mammals therefore becomes technologically difficult. However, the outlier approach based on empirical data is free from any assumption of demographic history, and is considered robust to avoid the potential confounding effect of population demographic history. Here, we employed an outlier approach based on genome-wide empirical data to retrieve the top 1% of genes showing high level *iFXD* values.

### Shared positive selection of genes among different species

One-to-one orthologous genes between dogs and pigs, horses, cattle and sheep were downloaded from Ensembl (http://www.ensembl.org/) using BioMart. One-to-one orthologous genes between dogs and goats were retrieved by a reciprocal best-hit BLASTP search. Overall, 10,103 one-to-one orthologous genes were identified among dogs, pigs, horses, cattle, sheep and goats. The value showing convergent signature of positive selection for each gene among the six mammals was identified as 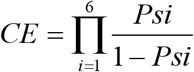, where *i* represents the different mammals and *P*s represent the probability of genes being under positive selection in the mammal. In total, 30 candidate genes were identified to harbor a high *iFXD* value (95% percentile rank) in at least three species.

### Data simulation

Simulation was performed by randomly choosing the same number of genes as the number of positively selected genes in each species. The number of shared genes was counted among six simulated data. 1000 random sampling were performed in total.

### Gene-gene interaction data

Protein-protein interactions were compiled from two resources, InWeb (InWeb3_HC_NonRed) (Rossin et al. 2011) and BioGRID (BIOGRID-ALL-3.3.124.tab) (Stark et al. 2006). We used only non-redundant interactions, and defined all interactions as undirected edges in a binary network. The ratio of gene-gene interaction of a group of genes, or between two groups of genes, was defined as the number of gene-gene interactions divided by the total number of gene pairs. Namely, within a group of gene, R=I/(N*(N-1)), and between two groups of genes, R=I/(N1*N2), where R is the ratio, I is the number of gene-gene interactions, and N, N1 and N2 are the numbers of genes. Simulation was performed by randomly choosing genes with the same number as positively selected genes in each species. 1000 simulations were performed, and the simulated ratios of gene-gene interactions were averaged.

### Association of variant at gene *EPAS1* with physiological parameters In Tibetan horse

As described in our pervious study (Gou et al. 2014), the foreleg venous blood was collected to measure physiological parameters in situ. 21 hematologic and hemorheologic parameters were measured by BC-2800Vet Auto Hematology Analyzer (Mindray Co., Ltd, Shenzhen) and ZL1000 Auto Blood Rheology Analyzer (Zonci Co., Ltd, Beijing), respectively. Mutations in *EPAS1* were genotyped by Sanger sequencing technology. The *F* test in the analysis of variance (ANOVA) was used to evaluate the significance of genotypes. Association was performed in horses at the same elevation.

### Transcriptomic Analysis of Lung Tissues of Tibetan Pigs and Min pigs

Four adult (6 months old) animals from each of the Tibetan and Min breeds were used for this analysis. For each breed, two females (not pregnant) and two male individuals were sampled. The Tibetan pigs were collected from Hezuo city in Gannan Tibet Autonomous Prefecture of Gansu Province (highland), and the Min pigs were sampled from Gongzhuling in Jilin Province (lowland). Animals were slaughtered and lung tissue immediately frozen in liquid nitrogen after collection and then stored at −80°C until RNA extraction. Total RNA was isolated from each tissue using TRlzol (Invitrogen) following the manufacturer’s protocols.

Sequencing libraries were prepared using the TruSeq™ RNA Sample Preparation Kit (Illumina, San Diego, USA) according to the manufacturer’s instructions. RNAs with poly (A) tails were isolated from total RNA using oligo (dT) magnetic beads, and was further fragmented. The first-strand cDNA was synthesized using reverse transcriptase and random hexamer primers, and RNase H and DNA polymerase I were used to synthesize the second-strand cDNA. The cDNA library had an insert size of 300-400 base pair (bp), and paired-end library was sequenced on Illumina Hiseq 2000 sequencing platform with a read length of 100 bp.

Raw reads were trimmed to remove adaptor sequences, low quality bases (Phred quality score <20) at the end of the RNA-seq reads, and subsequently filtered for length >25 bp using Trimmomatic [*Trimmomatic: a flexible trimmer for Illumina sequence data*]. The trimmed reads were then aligned against the reference genome of *Sus scrofa* (Sscrfa 10.2) using TopHat v2.0.4 with default parameters (Trapnell et al. 2009) and using the gene annotation available at Ensembl v77 (ftp://ftp.ensembl.org/pub/release-77/gtf/sus_scrofa). Transcripts were first assembled using the reference annotation based transcript (RABT) assembly method using Cufflinks (Trapnell et al. 2010). Novel transcripts were subsequently extracted from the output of the assemblers using a custom Perl script. The reference annotation was then merged with the novel transcripts to generate a new reference annotation file. To quantify gene expression, the FPKM (Fragments Per Kilobase of transcript per Million mapped reads) values were calculated using Cuffmerge without RABT. Finally, CuffDiff was applied to identify differentially expressed genes (Trapnell et al. 2010). We considered those with FDR (false discovery rate) <0.01 as significantly differentially expressed genes between high- and low-land pig breeds. In addition, DESeq (Anders and Huber 2012) and edgeR (Robinson et al. 2010) programs were used to identify differentially expressed genes.

### Experiments to study the function of *C10orf67*

To study the function of *C10orf67*, independent shRNAs targeting different regions of the gene were constructed using the pLKO.1 vector to knockdown *C10orf67* expression in NIH-3T3 cells. The 21 bp scrambled shRNA sequences were GCACTACCAGAGCTAACTCAG and ACCCTCTGAGTTAACATTTAA; the *mC10orf67* shRNA#1 sequence was GGGCAGAGAATGAAAGGTTAA; and the *mC10orf67* shRNA#2 sequence was AGCTTTCATGTCCTAAAGAAT. The sequences of all constructs were verified by Sanger sequencing. Lentiviruses were generated according to the manufacturer’s protocol. In brief, supernatants containing different lentiviruses generated from HEK-293T cells were collected at 48 and 72 h post-transfection, respectively; cells were infected successively twice by 48 h and 72 h viruses in the presence of 4 μg/mL polybrene. Stable cell lines were selected using puromycin. For the cell proliferation assay, tested cell lines were plated into 24-well plates and cell numbers were counted each day. The numbers of apoptotic cells were determined by flow cytometry using FITC-conjugated annexin V (BD Pharmingen) staining. The pipeline of Tophat, Cufflinks and Cuffdiff was employed to find the differentially expressed genes between *C10orf67* knockdown and scrambled-shRNA control NIH-3T3 cells under normoxia and hypoxia. Gene enrichment analysis was performed by g:profiler (Reimand et al. 2011).

### Gene enrichment analysis

We employed g:profiler (Reimand et al. 2011) (http://biit.cs.ut.ee/gprofiler) for enrichment analysis with Benjamini-Hochberg FDR as the multiple correction. We look for enriched modules in BioGRID protein–protein interaction (PPI) network by g:profiler program. For each species, only one significant gene-gene interaction network was found (P<0.01), and the network contains gene *EPAS1*.

## Data availability

The genome raw data is available from the SRA under bioproject number PRJNA281979.

## Acknowledgements

DDW, YBC were supported by the Strategic Priority Research Program of the Chinese Academy of Sciences (XDB13000000), National Natural Science Foundation of China (91731304, 31321002) and the Chinese 973 program (2013CB835200, 2013CB835204), YBC was also supported by grants from the National Natural Science Foundation of China (81322030, 31271579), Yunnan Province High-level Talents Introduced Program and Yunnan Applied Basic Research Projects (2014FA038). XG was supported by Chinese 863 program (2013AA102503), and the National Natural Science Foundation of China (31260542, 31260543, 31160449, 31460581, 31560617). YHM was supported by the National Natural Science Foundation of China (31272403). DDW was supported by the Youth Innovation Promotion Association, Chinese Academy of Sciences.

