## Supplementary Materials for "Convergent genomic signatures of high altitude adaptation among domestic mammals"

### Supplementary Note:

#### Inference of demographic history and population simulation

We first filtered variants that were missing with >50%. The genotypes were then imputed by Beagle (Browning et al. 2018). Samples of each species were classified as two populations: highland population and lowland population. To eliminate the uncertainty of ancestral state of each allele, we extracted the joint folded SFS (Li and Stephan 2006; Gutenkunst et al. 2009) based on imputed genotypes. We calculated the likelihood function for different demographic models by fastsimcoal2 (Excoffier et al. 2013). For each model, 200 to 300 runs were carried out. 100000 coalescent simulations per likelihood estimation and 20 to 40 expectation-conditional maximization (ECM) cycles were used for each run. The Akaike information criterion (AIC) (Akaike 1974) was used to compare different models.

As there are too many demographic models to be compared, it is difficult to infer the demographic history of two populations simultaneously. Therefore, we used an Ancestral-to-Derived Hierarchical Search strategy (Zeng et al. 2018), which assumes that the derived population does not influence the demography of the ancestral population. Because highland domestic animals migrated from lowland populations, we first compared three possible demographic models of lowland population for each species (Supplementary Figure 1; Supplementary Table 5). The bottleneck model is the best demographic model for all six species. To further explore how highland population migrated to Tibetan Plateau, another two possible joint demographic models were taken into consideration given that highland population was supposed to have experienced a bottleneck when migrating to Tibetan Plateau (Supplementary Figure 2; Supplementary Tables 6-7). Because we focused on recent demographic events, we referred to domestic time and generation interval of these species when setting parameters (Supplementary Tables 8-9).

To make sure the accuracy of our method, 1000 simulation sequence data (upstream 1Mb + 50kb + down 1Mb) were generated under best demographic model for each species using fastsimcoal2. The $F_{ST}$ (Akey et al. 2002) of each SNP was calculated as previously described. $F_{ST}$ with negative values having no biological explanation and were arbitrarily set to 0. selscan (Szpiech and Hernandez 2014) was used to calculate the XP-EHH (Sabeti et al. 2007) value for each SNP. ΔDAF was calculated for each SNP as the DAF in the domestic animals from the highlands minus the DAF in domestic animals from the lowlands. When compared to simulation data, all these candidate positively selected genes are still outlier compared to simulation data based on *iFXD* value. These positively selected genes based on the outlier approach still exhibit extreme divergence based on $F_{ST}$, ΔDAF and XPEHH values (Supplementary Figure 4). It indicates that our outlier approach is robust to demographic history.

To evaluate the influence of SNP density, we perform further analysis to choose the SNP harboring the highest *iFXD* value within a gene to represent the value of this gene. For each species, most of candidate positively selected genes by the two different strategies are overlapped (139 in sheep, 139 in dog, 146 in horse, 169 in goat, 155 in cattle, and 130 in pig). It suggests that the candidate positively selected genes detected by our outlier method are robust to SNP density.

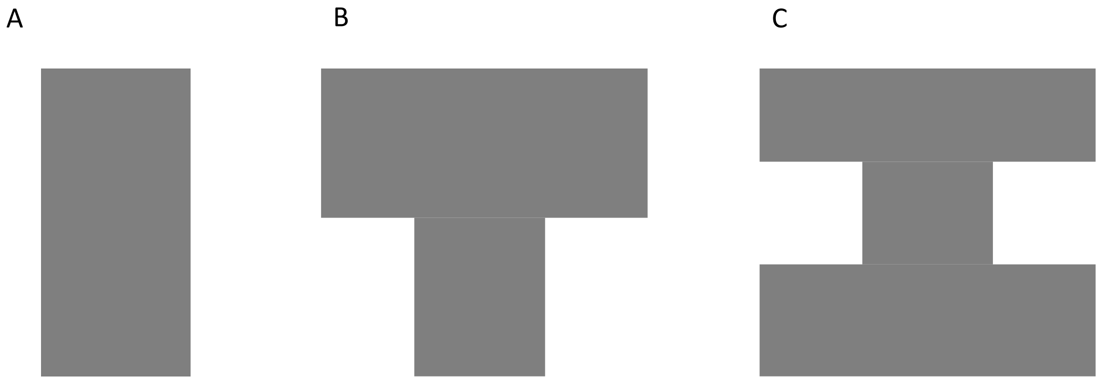


Supplementary Figure S1: three possible demographic models of lowland population for each species.

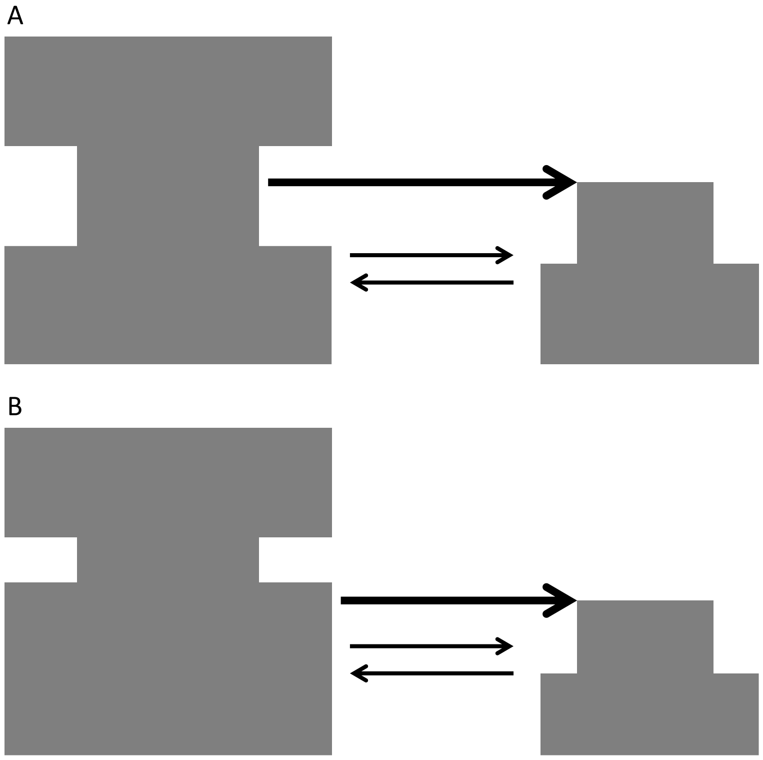


Supplementary Figure S2: two possible joint demographic models were taken into consideration given that highland population was supposed to have experienced a bottleneck when migrating to Tibetan Plateau.

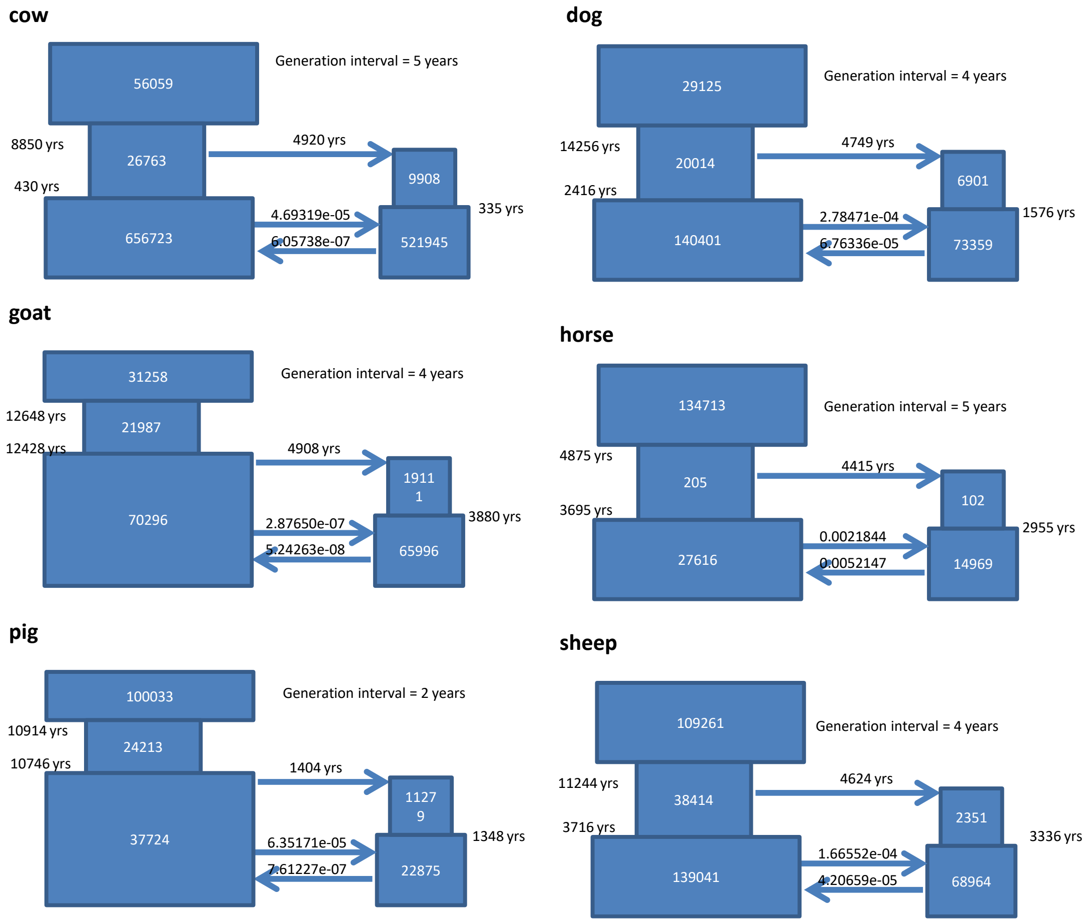


Supplementary Figure S3: Models of demographic history inferred by fastsimcoal2 program.

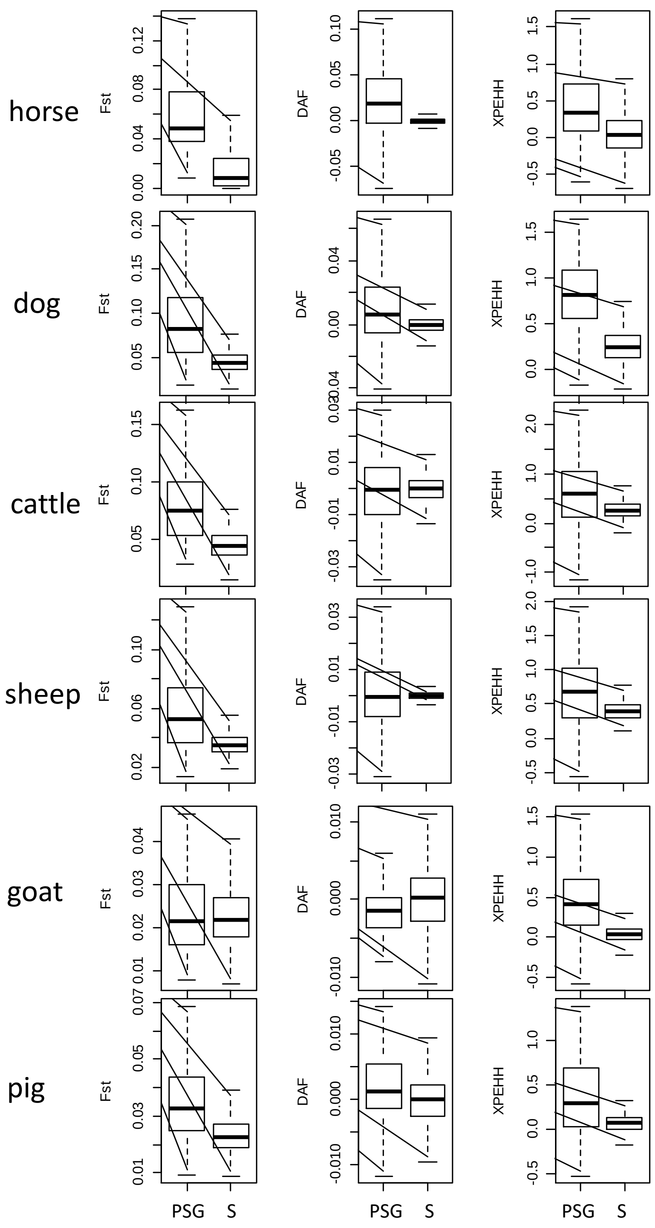


Supplementary Figure S4: Comparison of Fst, ΔDAF and XPEHH values of candidate positively selected genes (PSG) with simulated data (S) reveal extreme divergence of candidate positively selected genes.

| Gene | dog | horse | pig | cattle | sheep | goat |
|---|---|---|---|---|---|---|
| *EPAS1* | 1 | 1 | 1 | 0 | 1 | 1 |
| *C10orf67* | 1 | 0 | 1 | 1 | 0 | 0 |
| *ACADVL* | 1 | 0 | 1 | 0 | 0 | 0 |
| *AFF3* | 0 | 0 | 1 | 0 | 0 | 1 |
| *ANKRD6* | 1 | 0 | 0 | 1 | 0 | 0 |
| *ARHGAP26* | 1 | 0 | 0 | 0 | 1 | 0 |
| *ATAD1* | 0 | 1 | 0 | 0 | 0 | 1 |
| *ATP10A* | 0 | 0 | 0 | 1 | 1 | 0 |
| *BRDT* | 0 | 0 | 0 | 1 | 0 | 0 |
| *C2orf80* | 0 | 1 | 1 | 0 | 0 | 0 |
| *DLAT* | 0 | 0 | 0 | 1 | 1 | 0 |
| *DSG3* | 0 | 0 | 1 | 0 | 0 | 1 |
| *DYSF* | 1 | 0 | 1 | 0 | 0 | 0 |
| *DZIP3* | 0 | 0 | 1 | 0 | 0 | 1 |
| *ESRRG* | 0 | 0 | 1 | 0 | 0 | 1 |
| *FERMT2* | 0 | 0 | 0 | 1 | 1 | 0 |
| *FGF5* | 1 | 0 | 0 | 0 | 0 | 1 |
| *GAB1* | 0 | 0 | 0 | 1 | 0 | 1 |
| *GPBP1* | 1 | 0 | 0 | 1 | 0 | 0 |
| *GPCPD1* | 0 | 0 | 0 | 1 | 1 | 0 |
| *GULO* | 1 | 1 | 0 | 0 | 0 | 0 |
| *HCN4* | 0 | 1 | 0 | 1 | 0 | 0 |
| *IKZF1* | 0 | 0 | 1 | 0 | 1 | 0 |
| *JAZF1* | 0 | 0 | 0 | 0 | 1 | 1 |
| *LRRFIP1* | 0 | 0 | 0 | 0 | 1 | 1 |
| *LTF* | 0 | 0 | 0 | 0 | 1 | 1 |
| *NELL1* | 1 | 0 | 0 | 0 | 0 | 1 |
| *OR51B4* | 1 | 1 | 0 | 0 | 0 | 0 |
| *PAPSS2* | 0 | 1 | 0 | 0 | 0 | 1 |
| *PCNP* | 0 | 1 | 1 | 0 | 0 | 0 |
| *PDE4B* | 0 | 0 | 0 | 1 | 1 | 0 |
| *PELP1* | 1 | 0 | 1 | 0 | 0 | 0 |
| *PPP1R8* | 1 | 0 | 0 | 1 | 0 | 0 |
| *SAFB* | 0 | 0 | 0 | 0 | 1 | 0 |
| *TBC1D31* | 0 | 0 | 1 | 0 | 1 | 0 |
| *TEX14* | 0 | 1 | 0 | 0 | 1 | 0 |
| *TNKS1BP1* | 1 | 0 | 0 | 0 | 1 | 0 |
| *TXNDC16* | 0 | 1 | 0 | 1 | 0 | 0 |
| *UPB1* | 0 | 1 | 1 | 0 | 0 | 0 |
| *UTRN* | 0 | 0 | 0 | 1 | 0 | 1 |
| *ZFAND3* | 1 | 1 | 0 | 0 | 0 | 0 |
| *ZNF197* | 1 | 1 | 0 | 0 | 0 | 0 |
| *ZNF35* | 1 | 1 | 0 | 0 | 0 | 0 |

**Supplementary Figure S5: Convergent positive selection of 43 genes among the six domestic mammals.** 1 indicates that the gene evolved under positive selection as it was in the top 1% ranking.

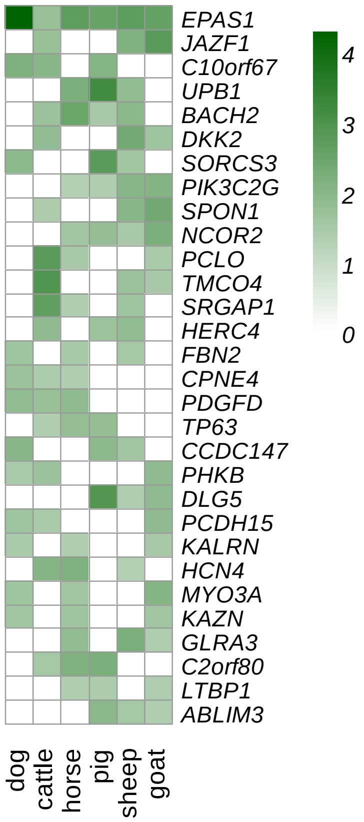


**Supplementary Figure S6: Convergent positive selection of genes among the six domestic mammals.** Heatmap of 30 genes evolving under positive selection in at least three species. White represents no significance (*P*>0.05). The other colors in the heatmap represent negative log10 transformed *P*-values of genes under positive selection.

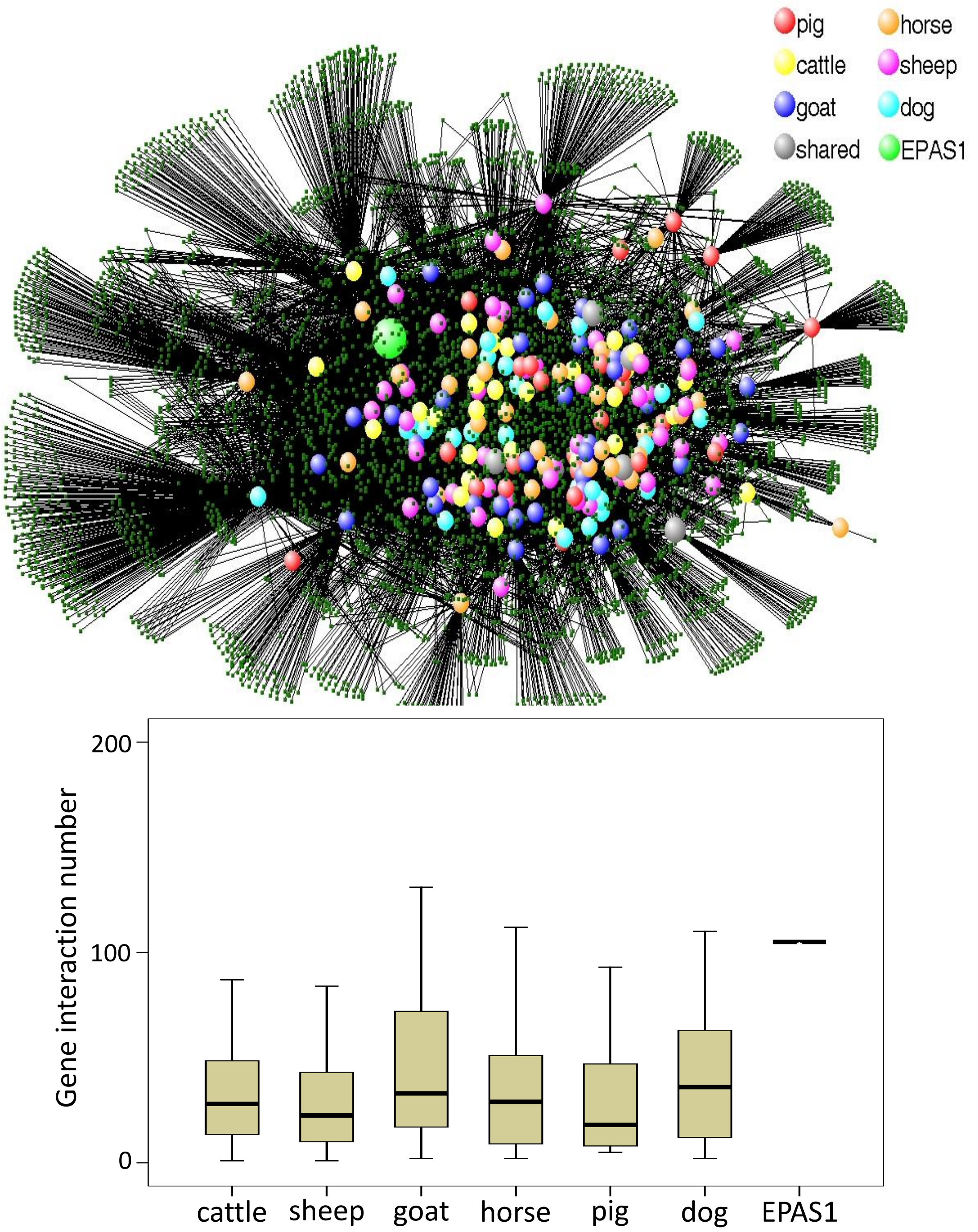


**Supplementary Figure S7: Gene-gene interaction network enriched within candidate positively selected genes in six Tibetan domestic mammals.** *EPAS1* was considered to be under positive selection in Tibetan cattle at the 95% percentile rank, but at the 99% percentile rank in other Tibetan mammals. Above is the interaction network, while below represents gene interaction numbers of positively selected genes in the above network.

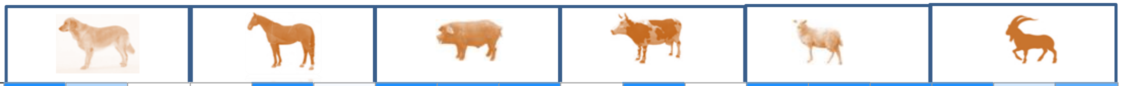

| | | | | | | | | | | | | | | | | | | | |
|---|---|---|---|---|---|---|---|---|---|---|---|---|---|---|---|---|---|---|---|
| response to hypoxia | GO:0001666 | 1.00e+00 | 3.51e-01 | 9.15e-22 | 2.39e-33 | 1.00e+00 | 3.20e-02 | 1.00e+00 | 9.43e-01 | 1.00e+00 | 0.00e+00 | 1.00e+00 | 1.24e-44 | 1.00e+00 | 1.00e+00 | 1.00e+00 | 1.00e+00 | 2.17e-01 | 7.22e-01 |
| regulation of transcription from RNA polymerase II promoter in response to hypoxia | GO:0061418 | 9.81e-85 | 7.69e-02 | 5.50e-52 | 1.91e-02 | 6.57e-02 | 9.91e-01 | 0.00e+00 | 9.81e-01 | 1.00e+00 | 1.00e+00 | 9.76e-01 | 8.85e-11 | 1.00e+00 | 1.00e+00 | 1.00e+00 | 8.71e-01 | 1.00e+00 | 1.00e+00 |
| positive regulation of transcription from RNA polymerase II promoter in response to hypoxia | GO:0061419 | 1.00e+00 | 8.05e-01 | 8.63e-30 | 1.00e+00 | 7.49e-02 | 5.83e-01 | 3.75e-54 | 7.99e-01 | 1.00e+00 | 1.83e-01 | 7.72e-09 | 6.85e-02 | 1.00e+00 | 1.00e+00 | 1.00e+00 | 1.36e-10 | 1.00e+00 | 5.83e-02 |
| negative regulation of transcription from RNA polymerase II promoter in response to hypoxia | GO:0061428 | 1.00e+00 | 2.56e-01 | 3.64e-01 | 9.99e-01 | 8.81e-01 | 1.38e-09 | 1.73e-30 | 2.60e-01 | 9.99e-01 | 1.00e+00 | 1.00e+00 | 2.64e-43 | 1.31e-35 | 8.76e-36 | 1.18e-35 | 1.00e+00 | 9.94e-01 | 2.10e-02 |
| detection of hypoxia | GO:0070483 | 1.26e-18 | 2.33e-01 | 1.21e-02 | 1.00e+00 | 8.47e-01 | 1.79e-04 | 1.99e-03 | 6.43e-01 | 1.41e-15 | 1.22e-04 | 1.46e-05 | 1.00e+00 | 7.10e-34 | 3.78e-37 | 6.36e-34 | 1.97e-25 | 1.22e-01 | 1.00e+00 |
| cellular response to hypoxia | GO:0071456 | 1.00e+00 | 3.99e-07 | 5.89e-144 | 1.00e+00 | 7.67e-03 | 3.41e-06 | 0.00e+00 | 8.16e-01 | 3.08e-04 | 1.00e+00 | 1.00e+00 | 5.52e-110 | 4.44e-132 | 2.99e-117 | 7.62e-131 | 9.79e-01 | 1.00e+00 | 1.00e+00 |
| hypoxia-inducible factor-1alpha signaling pathway | GO:0097411 | 1.00e+00 | 1.00e+00 | 1.00e+00 | 1.00e+00 | 2.56e-01 | 1.00e+00 | 1.00e+00 | 2.77e-01 | 9.88e-01 | 9.41e-01 | 7.46e-02 | 1.00e+00 | 1.00e+00 | 1.00e+00 | 1.00e+00 | 1.13e-33 | 8.60e-01 | 7.41e-01 |
| regulation of cellular response to hypoxia | GO:1900037 | 9.99e-01 | 7.62e-03 | 1.00e+00 | 8.51e-07 | 9.03e-01 | 5.51e-01 | 1.00e+00 | 7.25e-01 | 7.53e-01 | 9.37e-11 | 9.81e-01 | 8.19e-03 | 1.00e+00 | 1.00e+00 | 1.00e+00 | 6.80e-124 | 1.00e+00 | 1.14e-05 |
| negative regulation of cellular response to hypoxia | GO:1900038 | 2.57e-45 | 8.13e-01 | 9.59e-01 | 4.80e-36 | 9.99e-01 | 8.79e-01 | 1.45e-62 | 9.67e-01 | 6.99e-09 | 1.00e+00 | 1.00e-02 | 1.00e+00 | 1.00e+00 | 1.00e+00 | 1.00e+00 | 1.00e+00 | 5.15e-01 | 3.52e-03 |
| positive regulation of hypoxia-inducible factor-1alpha signaling pathway | GO:1902073 | 7.09e-19 | 4.06e-01 | 1.00e+00 | 1.00e+00 | 1.11e-04 | 1.53e-03 | 1.12e-02 | 5.95e-01 | 4.64e-01 | 2.09e-01 | 3.07e-02 | 6.46e-01 | 3.97e-04 | 4.62e-04 | 3.88e-04 | 9.91e-01 | 8.13e-01 | 7.50e-01 |
| intrinsic apoptotic signaling pathway in response to hypoxia | GO:1990144 | 1.00e+00 | 3.61e-05 | 7.03e-04 | 1.00e+00 | 1.06e-01 | 1.55e-01 | 1.00e+00 | 6.75e-02 | 2.60e-05 | 1.61e-11 | 4.13e-02 | 8.97e-01 | 1.00e+00 | 1.00e+00 | 1.00e+00 | 1.00e+00 | 1.00e+00 | 1.00e+00 |
| positive regulation of proteasomal ubiquitin-dependent protein catabolic process involved in cellular response to hypoxia | GO:2000777 | 1.00e+00 | 9.62e-02 | 1.01e-03 | 4.83e-01 | 4.80e-01 | 1.72e-02 | 3.30e-18 | 1.60e-01 | 9.79e-01 | 1.00e+00 | 1.00e+00 | 1.97e-57 | 3.50e-08 | 1.44e-08 | 3.37e-08 | 1.00e+00 | 3.60e-01 | 8.63e-01 |

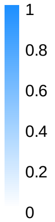


XP-EHH ΔDAF $F_{ST}$

**Supplementary Figure S8: Significances of XP-EHH, $F_{ST}$ and ΔDAF of SNPs within genes in hypoxia related gene ontology is higher than within other genes.**

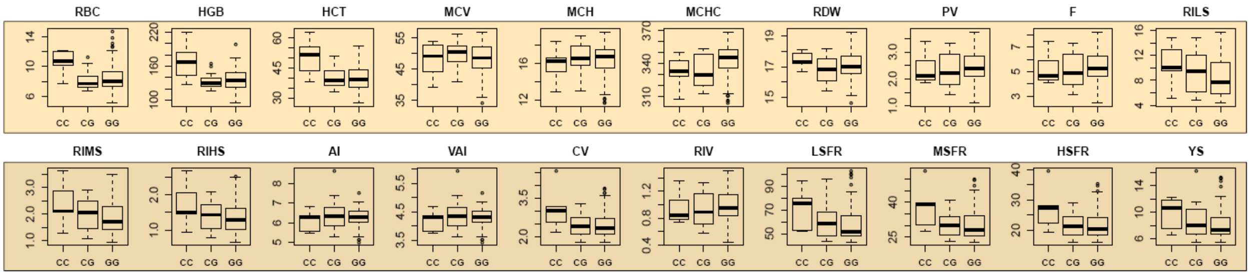


**Supplementary Figure S9: Association of SNP (chr15: 52623471) within horse *EPAS1* with 20 blood related traits.** The frequency of the derived allele (G) of this SNP is 0.7939 in the Tibetan horse, and 0.278 in the lowland horse. The SNP was found to be significantly associated with HGB and HCT ($P<0.01$, corrected by FDR). The numbers of horse with each genotype are CC: 7, GG: 75, GC: 16. RBC: Red blood cell (RBC) (10E6/ul), HGB: Hemoglobin (HGB) (g/L), HCT: Hematocrit (HCT) (%), MCV: Mean corpuscular volume (MCV) (fl), MCH: Mean corpsular hemoglobin (MCH) (pg), MCHC: Mean corpuscular hemoglobin concentration (MCHC) (g/L), RDW: Red blood cell distribution width (RDW) (%), PV: Plasma viscosity (mPa·s), F: Fibrinogen (g/L), RILS: Relative index of low shear, RIMS: Relative index of middle shear, RIHS: Relative index of high shear, AI: Erythrocyte aggregation index (AI), VAI: Viscometric aggregation index (VAI), CV: Casson viscosity (mPa·s), RIV: RBC internal viscosity (mPa·s), LSFR: Low shear flow resistance (10E9 SI), MSFR: Middle shear flow resistance (10E9 SI), HSFR: High shear flow resistance (10E9 SI), YS: Yield stress (mPa).

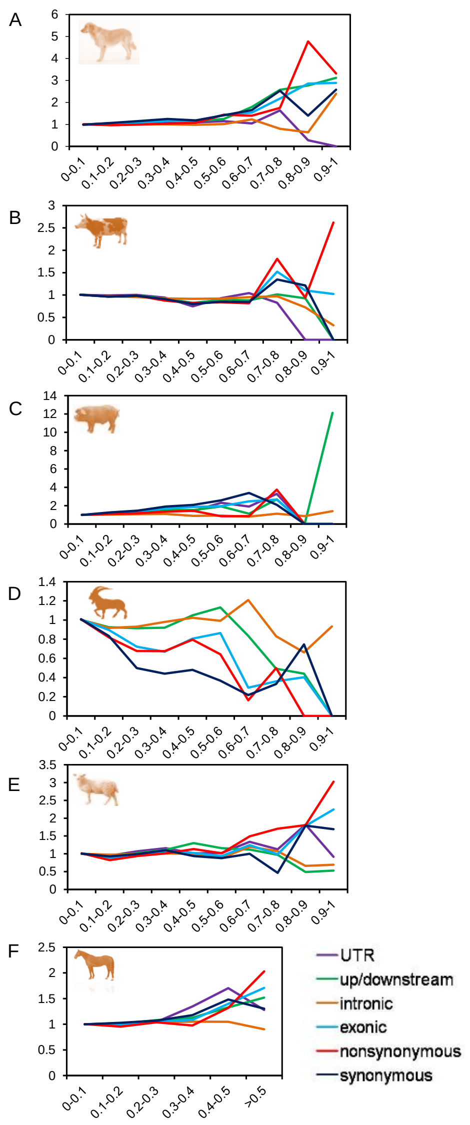


**Supplementary Figure S10: $F_{ST}$ of SNPs between domestic mammals from the Tibetan Plateau and lowlands.** Different types of SNPs were divided into 10 bins according to their $F_{ST}$ values. The enrichment level of SNPs in each $F_{ST}$ bin was calculated by the proportion of the SNPs divided by the proportion of intergenic SNPs.

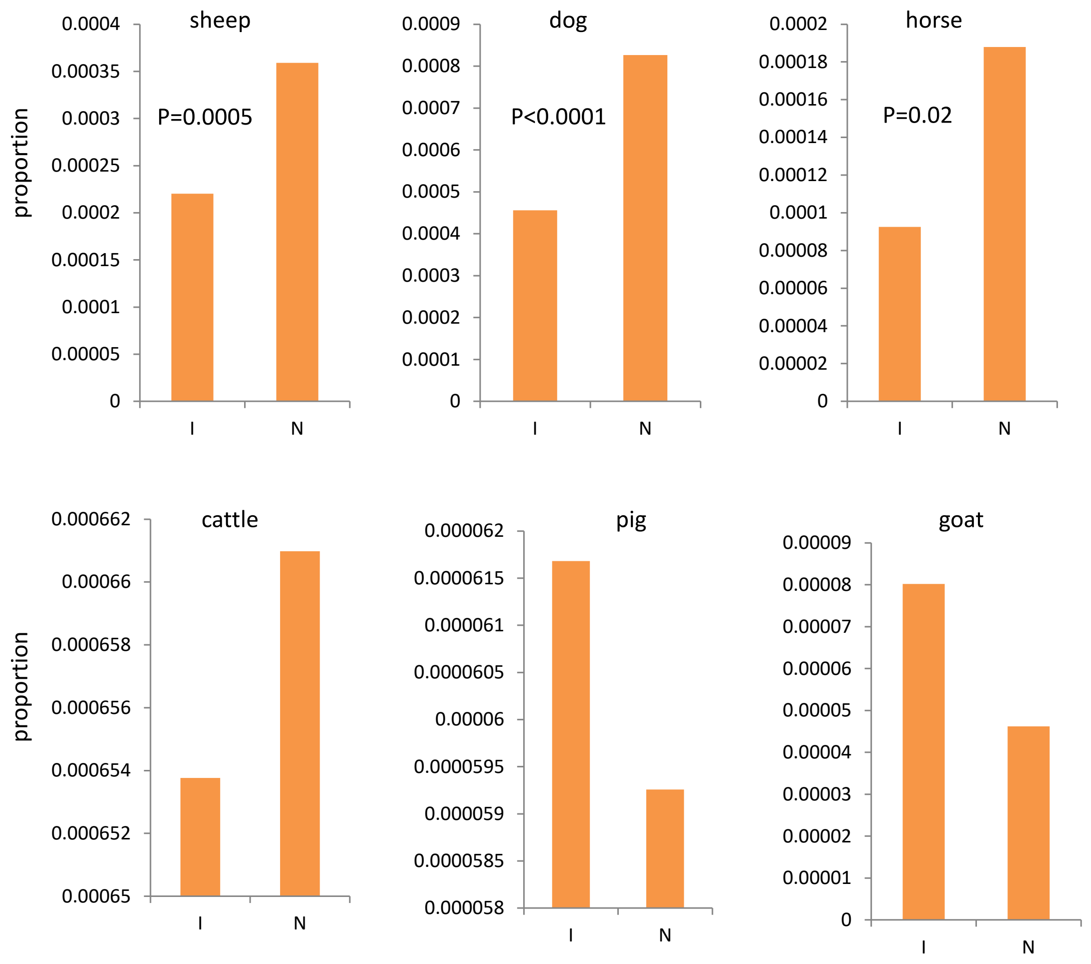


Supplementary Figure S11: Proportion of SNPs harboring higher Fst value (≥0.6) among intergenic SNPs (I) and non-synonymous SNPs (N) in different species. The statistical significance was calculated by $X^2$ test.

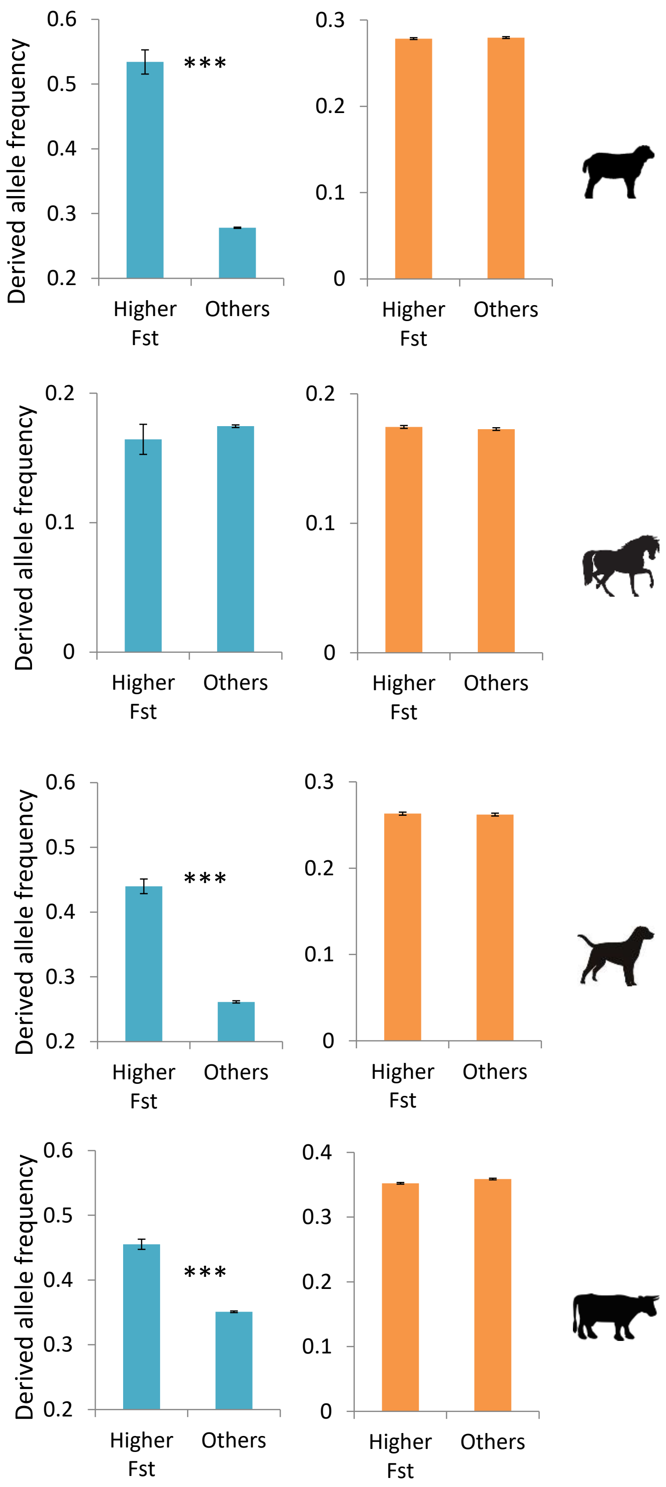


Supplementary Figure S12: Derived allele frequencies of non-synonymous SNPs harboring higher Fst value (top 1%) between highland and lowland populations and derived allele frequencies of other non-synonymous SNPs at highland population (left panel) and lowland population (right panel). *** P<0.001 by Mann-Whitney U test).

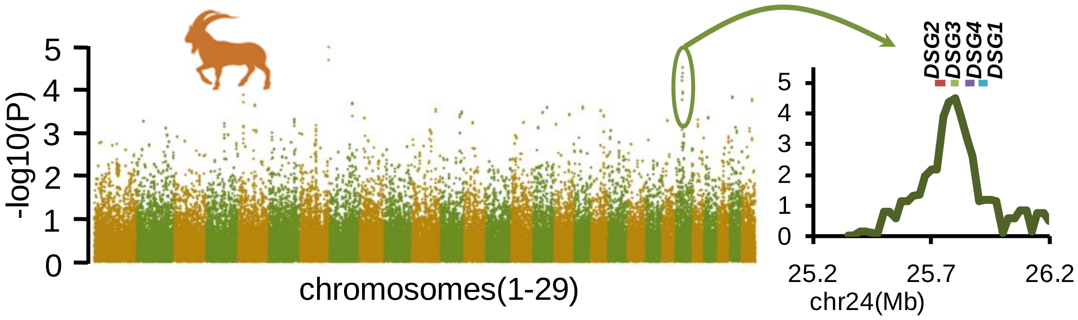


**Supplementary Figure S13: Genomic landscape of *iFXD* values in the genome of the Tibetan goat.** The horizontal axis represents 29 different autosomes. The ordinate presents the negative log transformed P-values of *iFXD* of each window. The signature of positive selection across the *DSG* gene cluster in chromosome 24 is presented on the right.

A

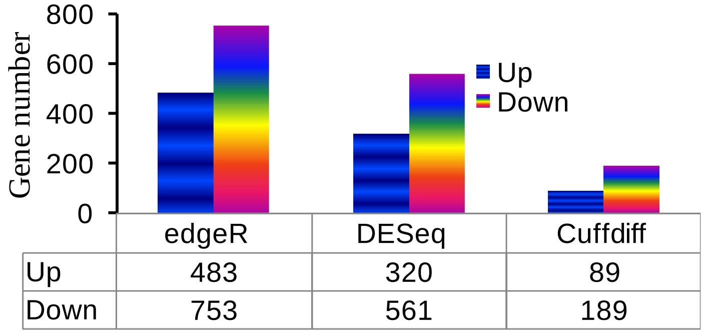


B

| GO Description | N | up | down | *P* |
|---|---|---|---|---|
| response to organic substance | 30 | 6 | 24 | 1.08E-05 |
| response to wounding | 24 | 9 | 15 | 1.24E-04 |
| extracellular matrix organization | 10 | 5 | 5 | 0.0067 |
| blood vessel development | 14 | 6 | 8 | 0.012 |
| inflammatory response | 16 | 6 | 10 | 0.013 |
| vasculature development | 14 | 6 | 8 | 0.016 |
| collagen fibril organization | 6 | 4 | 2 | 0.035 |

C

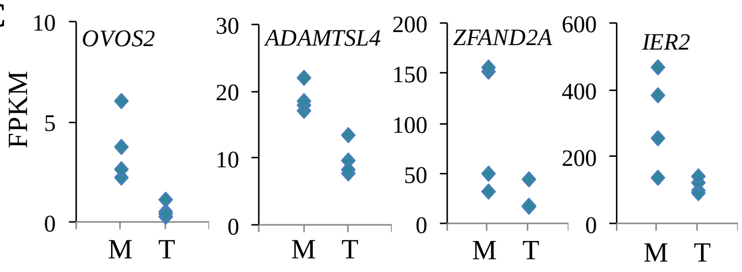


**Supplementary Figure S14: Genomic and transcriptomic analysis of positive selection in the Tibetan pig.** (A) A greater number of differentially expressed genes were down-regulated rather than up-regulated in the Tibetan pigs. (B) Categories enriched in the differentially expressed genes between lung transcriptomes of the Tibetan and Min pigs. *P* values are corrected by FDR implemented in DAVID program (Dennis et al. 2003). (C) Expression values (FPKM) of four positively selected genes in four Tibetan pigs (T) and four Min pigs (M).

**Supplementary Table S1: Genomes used in this study**. Shaded individuals were sequenced for this study.

| Breeds | Sample ID | Group | Source | Depth |
|---|---|---|---|---|
| Tibetan Horse | Yp9586 | Highland | QingHai | 6.21 |
| Tibetan Horse | Yp9588 | Highland | QingHai | 4.99 |
| Tibetan Horse | Yp9589 | Highland | QingHai | 5.56 |
| Tibetan Horse | Yp9682 | Highland | QingHai | 5.17 |
| Tibetan Horse | Yp9689 | Highland | QingHai | 5.12 |
| Tibetan Horse | Yp9782 | Highland | QingHai | 9.59 |
| Tibetan Horse | m14 | Highland | Tibet | 6.12 |
| Tibetan Horse | m2 | Highland | QingHai | 6.09 |
| Tibetan Horse | m8 | Highland | Tibet | 6.29 |
| Tibetan Horse | m9 | Highland | Tibet | 8.72 |
| Tibetan Horse | yp9581 | Highland | QingHai | 6.31 |
| Tibetan Horse | yp9590 | Highland | QingHai | 5.16 |
| Tibetan Horse | yp9591 | Highland | QingHai | 11.6 |
| Tibetan Horse | yp9592 | Highland | QingHai | 6.67 |
| Tibetan Horse | yp9593 | Highland | QingHai | 10.06 |
| Tibetan Horse | yp9680 | Highland | QingHai | 6.47 |
| Tibetan Horse | yp9683 | Highland | QingHai | 7.4 |
| Tibetan Horse | yp9684 | Highland | QingHai | 6.94 |
| Tibetan Horse | yp9687 | Highland | QingHai | 6.7 |
| Tibetan Horse | yp9783 | Highland | QingHai | 6.93 |
| Tibetan Horse | yp9784 | Highland | QingHai | 6.09 |
| Tibetan Horse | yp9785 | Highland | QingHai | 6.17 |
| Tibetan Horse | yp9786 | Highland | QingHai | 5.94 |
| Tibetan Horse | yp9679 | Highland | QingHai | 6.04 |
| Horse | Yp9308 | Lowland | HeiBei | 5.05 |
| Horse | Yp9324 | Lowland | HeiBei | 4.71 |
| Horse | Yp9381 | Lowland | XinJiang | 4.91 |
| Horse | Yp9426 | Lowland | XinJiang | 9.13 |
| Horse | Yp9430 | Lowland | XinJiang | 4.21 |
| Horse | yp9286 | Lowland | HeiBei | 7.12 |
| Horse | yp9287 | Lowland | HeiBei | 5.88 |
| Horse | yp9288 | Lowland | HeiBei | 6.12 |
| Horse | yp9345 | Lowland | HeiBei | 6.08 |

| | | | | |
|---|---|---|---|---|
| Horse | yp9423 | Lowland | XinJiang | 6.72 |
| Equus przewalskii | SRR899957 | out group | PRJNA205517 | 5.82 |
| Tibetan cattle | 100 | highland | GongBuJiangDa,Tibet | 6.42 |
| Tibetan cattle | 105 | highland | GongBuJiangDa,Tibet | 9.27 |
| Tibetan cattle | 108 | highland | GongBuJiangDa,Tibet | 7.58 |
| Tibetan cattle | 110 | highland | GongBuJiangDa,Tibet | 4.66 |
| Tibetan cattle | 113 | highland | GongBuJiangDa,Tibet | 3.97 |
| Tibetan cattle | 114 | highland | GongBuJiangDa,Tibet | 4.41 |
| Tibetan cattle | 115 | highland | GongBuJiangDa,Tibet | 6.15 |
| Tibetan cattle | 116 | highland | GongBuJiangDa,Tibet | 6.49 |
| Tibetan cattle | 117 | highland | GongBuJiangDa,Tibet | 4.62 |
| Tibetan cattle | 118 | highland | GongBuJiangDa,Tibet | 7.5 |
| Tibetan cattle | 119 | highland | GongBuJiangDa,Tibet | 5.97 |
| Tibetan cattle | 121 | highland | GongBuJiangDa,Tibet | 5.54 |
| Tibetan cattle | 89 | highland | GongBuJiangDa,Tibet | 7.16 |
| Tibetan cattle | 92 | highland | GongBuJiangDa,Tibet | 6.8 |
| Tibetan cattle | 93 | highland | GongBuJiangDa,Tibet | 6.65 |
| Tibetan cattle | 95 | highland | GongBuJiangDa,Tibet | 7.47 |
| Tibetan cattle | 96 | highland | GongBuJiangDa,Tibet | 10.05 |
| Tibetan cattle | 98 | highland | GongBuJiangDa,Tibet | 6.51 |
| Tibetan cattle | 99 | highland | GongBuJiangDa,Tibet | 9.74 |
| Mishima-Ushi | SAMD00013606 | lowland | PRJDB2660 | 13.13 |
| Mishima-Ushi | SAMD00013607 | lowland | PRJDB2660 | 12.91 |
| Mishima-Ushi | SAMD00013608 | lowland | PRJDB2660 | 7.94 |
| Mishima-Ushi | SAMD00013609 | lowland | PRJDB2660 | 14.07 |
| Mishima-Ushi | SAMD00013610 | lowland | PRJDB2660 | 10.52 |
| Mishima-Ushi | SAMD00013611 | lowland | PRJDB2660 | 12.3 |
| Mishima-Ushi | SAMD00013612 | lowland | PRJDB2660 | 15.11 |
| Mishima-Ushi | SAMD00013613 | lowland | PRJDB2660 | 9.09 |
| Korean Native Cattle | SRR934415 | lowland | PRJNA210519 | 13.3 |
| Korean Native Cattle | SRR934416 | lowland | PRJNA210519 | 13.85 |
| Korean Native Cattle | SRR934417 | lowland | PRJNA210519 | 13.09 |
| Korean Native Cattle | SRR934418 | lowland | PRJNA210519 | 13.55 |
| Korean Native Cattle | SRR934419 | lowland | PRJNA210519 | 12.87 |
| Korean Native Cattle | SRR934420 | lowland | PRJNA210519 | 13.8 |
| Korean Native Cattle | SRR934432 | lowland | PRJNA210519 | 13.81 |

| | | | | |
|---|---|---|---|---|
| Korean Native Cattle | SRR934433 | lowland | PRJNA210519 | 12.61 |
| Korean Native Cattle | SRR934434 | lowland | PRJNA210519 | 12.77 |
| Korean Native Cattle | SRR934435 | lowland | PRJNA210519 | 11.79 |
| Korean Native Cattle | SRR934436 | lowland | PRJNA210519 | 11.66 |
| Korean Native Cattle | SRR934437 | lowland | PRJNA210519 | 12.08 |
| MangShi Cattle | ms1 | lowland | MangShi,Yunnan | 6.07 |
| MangShi Cattle | ms13 | lowland | MangShi,Yunnan | 4.73 |
| MangShi Cattle | ms17 | lowland | MangShi,Yunnan | 7.17 |
| MangShi Cattle | ms18 | lowland | MangShi,Yunnan | 4.84 |
| MangShi Cattle | ms2 | lowland | MangShi,Yunnan | 5.37 |
| MangShi Cattle | ms24 | lowland | MangShi,Yunnan | 6.13 |
| MangShi Cattle | ms25 | lowland | MangShi,Yunnan | 7.52 |
| MangShi Cattle | ms29 | lowland | MangShi,Yunnan | 6.57 |
| MangShi Cattle | ms3 | lowland | MangShi,Yunnan | 5.59 |
| MangShi Cattle | ms30 | lowland | MangShi,Yunnan | 5.53 |
| Bos grunniens | SRR962825 | out group | PRJNA217895 | 5.21 |
| Tibetan Mastiff | SRR1105792 | Highland | PRJNA233638 | 14.36 |
| Tibetan Mastiff | SRR1138360 | Highland | PRJNA233638 | 13.01 |
| Tibetan Mastiff | SRR1138361 | Highland | PRJNA233638 | 13.32 |
| Tibetan Mastiff | SRR1138362 | Highland | PRJNA233638 | 12.11 |
| Tibetan Mastiff | SRR1138363 | Highland | PRJNA233638 | 13.1 |
| Tibetan Mastiff | SRR1138364 | Highland | PRJNA233638 | 13.48 |
| Tibetan Mastiff | SRR1138365 | Highland | PRJNA233638 | 15.1 |
| Tibetan Mastiff | SRR1138367 | Highland | PRJNA233638 | 12.94 |
| Tibetan Mastiff | SRR1138368 | Highland | PRJNA233638 | 15.11 |
| Tibetan Mastiff | SRR1138369 | Highland | PRJNA233638 | 15.05 |
| Tibetan Mastiff | TM | Highland | PRJNA192935 | 12.84 |
| Village Dog | Dog1 | lowland | PRJNA232497 | 6.09 |
| Village Dog | Dog10 | lowland | PRJNA232497 | 6.71 |
| Village Dog | Dog11 | lowland | PRJNA232497 | 6.18 |
| Village Dog | Dog12 | lowland | PRJNA232497 | 6.63 |
| Village Dog | Dog13 | lowland | PRJNA232497 | 6.75 |
| Village Dog | Dog14 | lowland | PRJNA232497 | 6.67 |
| Village Dog | Dog15 | lowland | PRJNA232497 | 6.51 |
| Village Dog | Dog2 | lowland | PRJNA232497 | 6.62 |
| Village Dog | Dog3 | lowland | PRJNA232497 | 6.45 |

| | | | | |
|---|---|---|---|---|
| Village Dog | Dog4 | lowland | PRJNA232497 | 6.01 |
| Village Dog | Dog6 | lowland | PRJNA232497 | 6.62 |
| Village Dog | Dog7 | lowland | PRJNA232497 | 5.58 |
| Village Dog | Dog8 | lowland | PRJNA232497 | 6.43 |
| Village Dog | Dog9 | lowland | PRJNA232497 | 6.13 |
| KunMing Dog | SRR1134656 | lowland | PRJNA233638 | 11.15 |
| KunMing Dog | SRR1135309 | lowland | PRJNA233638 | 16.03 |
| KunMing Dog | SRR1135544 | lowland | PRJNA233638 | 14.91 |
| KunMing Dog | SRR1137077 | lowland | PRJNA233638 | 15.24 |
| KunMing Dog | SRR1137093 | lowland | PRJNA233638 | 14.98 |
| KunMing Dog | SRR1138308 | lowland | PRJNA233638 | 15.14 |
| KunMing Dog | SRR1138309 | lowland | PRJNA233638 | 16.03 |
| KunMing Dog | SRR1138310 | lowland | PRJNA233638 | 13.48 |
| KunMing Dog | SRR1138311 | lowland | PRJNA233638 | 13.83 |
| KunMing Dog | SRR1138312 | lowland | PRJNA233638 | 15.95 |
| YuanJiang Dog | SRR1138313 | lowland | PRJNA233638 | 15.15 |
| YuanJiang Dog | SRR1138314 | lowland | PRJNA233638 | 12.5 |
| YuanJiang Dog | SRR1138315 | lowland | PRJNA233638 | 12.05 |
| YuanJiang Dog | SRR1138316 | lowland | PRJNA233638 | 12.16 |
| YuanJiang Dog | SRR1138317 | lowland | PRJNA233638 | 11.67 |
| YuanJiang Dog | SRR1138318 | lowland | PRJNA233638 | 12.93 |
| YuanJiang Dog | SRR1138331 | lowland | PRJNA233638 | 12.28 |
| YuanJiang Dog | SRR1138332 | lowland | PRJNA233638 | 12.82 |
| YuanJiang Dog | SRR1138333 | lowland | PRJNA233638 | 13.46 |
| YuanJiang Dog | SRR1138334 | lowland | PRJNA233638 | 14.29 |
| Golden_jackal | Golden_jackal | outgroup | | 4.98 |
| Tibetan Pig | batch3_104 | Highland | LinZhi,Tibet | 3.09 |
| Tibetan Pig | batch3_108 | Highland | LinZhi,Tibet | 2.31 |
| Tibetan Pig | batch4_105 | Highland | LinZhi,Tibet | 5.47 |
| Tibetan Pig | batch6_167 | Highland | LinZhi,Tibet | 3.51 |
| Tibetan Pig | batch6_168 | Highland | LinZhi,Tibet | 3.81 |
| Tibetan Pig | kizbatch1_601 | Highland | LinZhi,Tibet | 3.82 |
| Tibetan Pig | kizbatch1_602 | Highland | LinZhi,Tibet | 3.68 |
| Tibetan Pig | kizbatch1_603 | Highland | LinZhi,Tibet | 3.41 |
| Tibetan Pig | kizbatch1_604 | Highland | LinZhi,Tibet | 2.16 |
| Tibetan Pig | kizbatch1_605 | Highland | LinZhi,Tibet | 2.68 |
| Tibetan Pig | kizbatch1_606 | Highland | LinZhi,Tibet | 3.46 |

| | | | | |
|---|---|---|---|---|
| Tibetan Pig | SRR652257 | Highland | PRJNA186497 | 4.42 |
| Tibetan Pig | SRR652258 | Highland | PRJNA186497 | 4.24 |
| Tibetan Pig | SRR652259 | Highland | PRJNA186497 | 3.82 |
| Tibetan Pig | SRR652260 | Highland | PRJNA186497 | 4.97 |
| Tibetan Pig | SRR652261 | Highland | PRJNA186497 | 3.96 |
| Tibetan Pig | SRR652262 | Highland | PRJNA186497 | 5.39 |
| Tibetan Pig | SRR652263 | Highland | PRJNA186497 | 4.18 |
| Tibetan Pig | SRR652264 | Highland | PRJNA186497 | 4.27 |
| Tibetan Pig | SRR652265 | Highland | PRJNA186497 | 6.05 |
| Tibetan Pig | SRR652266 | Highland | PRJNA186497 | 4.12 |
| Tibetan Pig | SRR652267 | Highland | PRJNA186497 | 3.58 |
| Tibetan Pig | SRR652268 | Highland | PRJNA186497 | 6.46 |
| Tibetan Pig | SRR652269 | Highland | PRJNA186497 | 4.71 |
| Tibetan Pig | SRR652270 | Highland | PRJNA186497 | 4.25 |
| Tibetan Pig | SRR652302 | Highland | PRJNA186497 | 5.97 |
| Tibetan Pig | SRR652303 | Highland | PRJNA186497 | 4.98 |
| Tibetan Pig | SRR652304 | Highland | PRJNA186497 | 4.18 |
| Tibetan Pig | SRR652305 | Highland | PRJNA186497 | 5.19 |
| Tibetan Pig | SRR652306 | Highland | PRJNA186497 | 4.39 |
| Tibetan Pig | SRR652307 | Highland | PRJNA186497 | 5 |
| Tibetan Pig | SRR652327 | Highland | PRJNA186497 | 5.24 |
| Tibetan Pig | SRR652339 | Highland | PRJNA186497 | 4.21 |
| Tibetan Pig | SRR652340 | Highland | PRJNA186497 | 4.2 |
| Tibetan Pig | SRR652341 | Highland | PRJNA186497 | 4.56 |
| Tibetan Pig | SRR652342 | Highland | PRJNA186497 | 4.27 |
| Tibetan Pig | SRR652343 | Highland | PRJNA186497 | 3.96 |
| Tibetan Pig | SRR652344 | Highland | PRJNA186497 | 5.92 |
| Tibetan Pig | SRR652345 | Highland | PRJNA186497 | 4.94 |
| Tibetan Pig | SRR652346 | Highland | PRJNA186497 | 5.83 |
| Tibetan Pig | SRR652347 | Highland | PRJNA186497 | 4.93 |
| Jiangquhai | ERR173179 | Lowland | PRJEB1683 | 9.06 |
| Jiangquhai | ERR173199 | Lowland | PRJEB1683 | 7.76 |
| Meishan | ERR173200 | Lowland | PRJEB1683 | 7.66 |
| Meishan | ERR173201 | Lowland | PRJEB1683 | 7.03 |
| Meishan | ERR173202 | Lowland | PRJEB1683 | 8.96 |
| Wild Boar South China | ERR173219 | Lowland | PRJEB1683 | 4.93 |
| Wild Boar South China | ERR173220 | Lowland | PRJEB1683 | 8.79 |
| Wild Boar North China | ERR173221 | Lowland | PRJEB1683 | 4.86 |

| | | | | |
|---|---|---|---|---|
| Wild Boar North China | ERR173222 | Lowland | PRJEB1683 | 8.51 |
| Xiang | ERR173223 | Lowland | PRJEB1683 | 7.44 |
| Xiang | ERR173224 | Lowland | PRJEB1683 | 7.25 |
| Penzhou pig-1 | SRR652348 | Lowland | PRJNA186497 | 4.4 |
| Penzhou pig-2 | SRR652349 | Lowland | PRJNA186497 | 4.27 |
| Penzhou pig-3 | SRR652350 | Lowland | PRJNA186497 | 4.66 |
| Wujin pig-1 | SRR652351 | Lowland | PRJNA186497 | 4.95 |
| Wujin pig-2 | SRR652352 | Lowland | PRJNA186497 | 4.72 |
| Wujin pig-3 | SRR652353 | Lowland | PRJNA186497 | 4.2 |
| Ya'nan pig-1 | SRR652354 | Lowland | PRJNA186497 | 4.06 |
| Ya'nan pig-2 | SRR652355 | Lowland | PRJNA186497 | 4.17 |
| Ya'nan pig-3 | SRR652356 | Lowland | PRJNA186497 | 4.82 |
| Neijiang pig-1 | SRR652357 | Lowland | PRJNA186497 | 4.92 |
| Neijiang pig-2 | SRR652362 | Lowland | PRJNA186497 | 5.65 |
| Neijiang pig-3 | SRR652363 | Lowland | PRJNA186497 | 4.32 |
| Jinhua pig-1 | SRR652374 | Lowland | PRJNA186497 | 4.35 |
| Jinhua pig-2 | SRR652375 | Lowland | PRJNA186497 | 4.48 |
| Jinhua pig-3 | SRR652376 | Lowland | PRJNA186497 | 3.89 |
| Wild boar-1 | SRR652377 | Lowland | PRJNA186497 | 4 |
| Wild boar-2 | SRR652378 | Lowland | PRJNA186497 | 5.18 |
| Wild boar-3 | SRR652379 | Lowland | PRJNA186497 | 5.08 |
| Warthog | ERR173203 | out group | PRJEB1683 | 9.99 |
| Tibetan Sheep | R11 | Highland | AnDuo,Tibet | 8.77 |
| Tibetan Sheep | R12 | Highland | AnDuo,Tibet | 9.41 |
| Tibetan Sheep | R13 | Highland | AnDuo,Tibet | 9.66 |
| Tibetan Sheep | R14 | Highland | AnDuo,Tibet | 8.77 |
| Tibetan Sheep | R15 | Highland | AnDuo,Tibet | 8.53 |
| Tibetan Sheep | R16 | Highland | AnDuo,Tibet | 9.71 |
| Tibetan Sheep | R17 | Highland | AnDuo,Tibet | 8.09 |
| Tibetan Sheep | R18 | Highland | AnDuo,Tibet | 9.53 |
| Tibetan Sheep | R19 | Highland | AnDuo,Tibet | 9.58 |
| Tibetan Sheep | R20 | Highland | AnDuo,Tibet | 8.06 |
| Tibetan Sheep | R27 | Highland | AnDuo,Tibet | 8.35 |
| Tibetan Sheep | R28 | Highland | AnDuo,Tibet | 8.38 |
| Tibetan Sheep | R29 | Highland | AnDuo,Tibet | 8 |
| Tibetan Sheep | R30 | Highland | AnDuo,Tibet | 9.64 |
| Tibetan Sheep | R35 | Highland | QuZiKa,Tibet | 6.41 |

| | | | | |
|---|---|---|---|---|
| Tibetan Sheep | R36 | Highland | QuZiKa,Tibet | 8.85 |
| Tibetan Sheep | R37 | Highland | QuZiKa,Tibet | 10.4 |
| Tibetan Sheep | R38 | Highland | QuZiKa,Tibet | 7.23 |
| Tibetan Sheep | R39 | Highland | QuSong,Tibet | 6.78 |
| Tibetan Sheep | R40 | Highland | QuSong,Tibet | 7.6 |
| Sheep | SRR501837 | Lowland | PRJNA160933 | 14.27 |
| Sheep | SRR501838 | Lowland | PRJNA160933 | 11.98 |
| Sheep | SRR501839 | Lowland | PRJNA160933 | 10.86 |
| Sheep | SRR501840 | Lowland | PRJNA160933 | 9.27 |
| Sheep | SRR501841 | Lowland | PRJNA160933 | 13.68 |
| Sheep | SRR501842 | Lowland | PRJNA160933 | 14.89 |
| Sheep | SRR501843 | Lowland | PRJNA160933 | 11.16 |
| Sheep | SRR501844 | Lowland | PRJNA160933 | 9.79 |
| Sheep | SRR501845 | Lowland | PRJNA160933 | 9.86 |
| Sheep | SRR501846 | Lowland | PRJNA160933 | 9.49 |
| Sheep | SRR501848 | Lowland | PRJNA160933 | 11.73 |
| Sheep | SRR501849 | Lowland | PRJNA160933 | 8.66 |
| Sheep | SRR501850 | Lowland | PRJNA160933 | 11.53 |
| Sheep | SRR501851 | Lowland | PRJNA160933 | 12.77 |
| Sheep | SRR501852 | Lowland | PRJNA160933 | 13.21 |
| Sheep | SRR501853 | Lowland | PRJNA160933 | 11.77 |
| Sheep | SRR501854 | Lowland | PRJNA160933 | 12.91 |
| Sheep | SRR501855 | Lowland | PRJNA160933 | 14.17 |
| Sheep | SRR501856 | Lowland | PRJNA160933 | 9.47 |
| Sheep | SRR501857 | Lowland | PRJNA160933 | 17.86 |
| Sheep | SRR501859 | Lowland | PRJNA160933 | 13.4 |
| Sheep | SRR501860 | Lowland | PRJNA160933 | 9.04 |
| Sheep | SRR501861 | Lowland | PRJNA160933 | 14.29 |
| Sheep | SRR501862 | Lowland | PRJNA160933 | 14.81 |
| Sheep | SRR501863 | Lowland | PRJNA160933 | 7 |
| Sheep | SRR501864 | Lowland | PRJNA160933 | 10.48 |
| Sheep | SRR501865 | Lowland | PRJNA160933 | 16.44 |
| Sheep | SRR501866 | Lowland | PRJNA160933 | 15.28 |
| Sheep | SRR501867 | Lowland | PRJNA160933 | 11.69 |
| Sheep | SRR501868 | Lowland | PRJNA160933 | 12.47 |
| Sheep | SRR501869 | Lowland | PRJNA160933 | 12.73 |

| | | | | |
|---|---|---|---|---|
| Sheep | SRR501870 | Lowland | PRJNA160933 | 9.4 |
| Sheep | SRR501871 | Lowland | PRJNA160933 | 12.9 |
| Sheep | SRR501872 | Lowland | PRJNA160933 | 14.34 |
| Sheep | SRR501873 | Lowland | PRJNA160933 | 14.04 |
| Sheep | SRR501874 | Lowland | PRJNA160933 | 8.08 |
| Sheep | SRR501875 | Lowland | PRJNA160933 | 13.33 |
| Sheep | SRR501876 | Lowland | PRJNA160933 | 11.83 |
| Sheep | SRR501877 | Lowland | PRJNA160933 | 5.64 |
| Sheep | SRR501878 | Lowland | PRJNA160933 | 11.57 |
| Sheep | SRR501879 | Lowland | PRJNA160933 | 10.62 |
| Sheep | SRR501880 | Lowland | PRJNA160933 | 13.15 |
| Sheep | SRR501881 | Lowland | PRJNA160933 | 13.11 |
| Sheep | SRR501882 | Lowland | PRJNA160933 | 11.16 |
| Sheep | SRR501883 | Lowland | PRJNA160933 | 14.15 |
| Sheep | SRR501884 | Lowland | PRJNA160933 | 15 |
| Sheep | SRR501885 | Lowland | PRJNA160933 | 12.41 |
| Sheep | SRR501886 | Lowland | PRJNA160933 | 13.23 |
| Sheep | SRR501887 | Lowland | PRJNA160933 | 12.54 |
| Sheep | SRR501888 | Lowland | PRJNA160933 | 10.99 |
| Sheep | SRR501889 | Lowland | PRJNA160933 | 13.53 |
| Sheep | SRR501890 | Lowland | PRJNA160933 | 14.03 |
| Sheep | SRR501891 | Lowland | PRJNA160933 | 12.87 |
| Sheep | SRR501892 | Lowland | PRJNA160933 | 12.47 |
| Sheep | SRR501893 | Lowland | PRJNA160933 | 10.3 |
| Sheep | SRR501894 | Lowland | PRJNA160933 | 14.05 |
| Sheep | SRR501896 | Lowland | PRJNA160933 | 4.58 |
| Sheep | SRR501899 | Lowland | PRJNA160933 | 13.51 |
| Sheep | SRR501900 | Lowland | PRJNA160933 | 11.7 |
| Sheep | SRR501901 | Lowland | PRJNA160933 | 12.66 |
| Sheep | SRR501902 | Lowland | PRJNA160933 | 14.67 |
| Sheep | SRR501903 | Lowland | PRJNA160933 | 12.81 |
| Sheep | SRR501904 | Lowland | PRJNA160933 | 18.11 |
| Sheep | SRR501905 | Lowland | PRJNA160933 | 11.51 |
| Sheep | SRR501906 | Lowland | PRJNA160933 | 11.19 |
| Sheep | SRR501907 | Lowland | PRJNA160933 | 14.65 |
| Sheep | SRR501908 | Lowland | PRJNA160933 | 14.29 |

| | | | | |
|---|---|---|---|---|
| Sheep | SRR501909 | Lowland | PRJNA160933 | 14.06 |
| Sheep | SRR501910 | Lowland | PRJNA160933 | 8.91 |
| Sheep | SRR501911 | Lowland | PRJNA160933 | 11.66 |
| Ovis dalli | SRR501847 | outgroup | PRJNA160934 | 11.13 |
| Tibetan Goat | R01 | Highland | QuSong,Tibet | 9.37 |
| Tibetan Goat | R02 | Highland | QuSong,Tibet | 10.86 |
| Tibetan Goat | R03 | Highland | QuSong,Tibet | 9.88 |
| Tibetan Goat | R04 | Highland | QuSong,Tibet | 10.18 |
| Tibetan Goat | R06 | Highland | AnDuo,Tibet | 10.24 |
| Tibetan Goat | R07 | Highland | AnDuo,Tibet | 10.22 |
| Tibetan Goat | R08 | Highland | AnDuo,Tibet | 9.85 |
| Tibetan Goat | R09 | Highland | AnDuo,Tibet | 8.78 |
| Tibetan Goat | R10 | Highland | QuSong,Tibet | 9.67 |
| Tibetan Goat | R21 | Highland | QuSong,Tibet | 9.19 |
| Tibetan Goat | R22 | Highland | QuSong,Tibet | 9.51 |
| Tibetan Goat | R23 | Highland | QuZiKa,Tibet | 11.48 |
| Tibetan Goat | R24 | Highland | QuZiKa,Tibet | 4.94 |
| Tibetan Goat | R25 | Highland | AnDuo,Tibet | 7.1 |
| Tibetan Goat | R26 | Highland | AnDuo,Tibet | 8.28 |
| Tibetan Goat | R31 | Highland | QuZiKa,Tibet | 9.08 |
| Tibetan Goat | R32 | Highland | QuZiKa,Tibet | 9.54 |
| Tibetan Goat | R33 | Highland | QuZiKa,Tibet | 9.39 |
| Tibetan Goat | R34 | Highland | QuZiKa,Tibet | 7.78 |
| Goat | CM22 | Lowland | ChengDu | 14.37 |
| Goat | CW26 | Lowland | HeBei | 9.74 |
| Rangeland | ERR318225 | Lowland | PRJEB4371 | 5.76 |
| Rangeland | ERR318226 | Lowland | PRJEB4371 | 5.54 |
| Rangeland | ERR318227 | Lowland | PRJEB4371 | 7.04 |
| Rangeland | ERR318228 | Lowland | PRJEB4371 | 6.84 |
| Rangeland | ERR318229 | Lowland | PRJEB4371 | 6.01 |
| Goat | JN42 | Lowland | ShanDong | 12.94 |
| Goat | LH21 | Lowland | GuangDong | 13.71 |
| Goat | M9 | Lowland | SiChuan | 14.1 |
| Goat | T1 | Lowland | HuNan | 11.61 |
| Goat | T11 | Lowland | HuBei | 14.21 |
| Goat | T15 | Lowland | ChongQing | 9.36 |

| | | | | |
|---|---|---|---|---|
| Goat | T18 | Lowland | ShanDong | 14.27 |
| Goat | T2 | Lowland | ShanDong | 13.02 |
| Goat | T3 | Lowland | ShanX1 | 7.69 |
| Goat | T5 | Lowland | XinJiang | 10.26 |
| Goat | T6 | Lowland | ShanX1 | 9.89 |
| Goat | T8 | Lowland | YunNan | 9.99 |
| Goat | W10 | Lowland | Inner Mongolia | 11.91 |
| Tibetan Sheep | R37 | Out group | QuZiKa,Tibet | 10.4 |

**Supplementary Table S2: SNPs shared between pairs of species**. SNPs in vcf file from each domestic mammal were coordinated to human genome (hg19) by Liftover program. SNPs mapped to the same position in human genome in two Tibetan species are considered to be shared SNPs.

| Species pair | Number of sharing SNPs |
|---|---|
| dog cattle | 118860 |
| dog_goat | 131905 |
| dog_horse | 88390 |
| dog_pig | 188310 |
| dog_sheep | 69770 |
| goat-cattle | 400783 |
| goat-pig | 369624 |
| horse_cattle | 138145 |
| horse_goat | 149707 |
| horse_pig | 214675 |
| horse_sheep | 78929 |
| pig_cattle | 328518 |
| sheep_cattle | 127527 |
| sheep_goat | 147315 |
| sheep-pig | 176299 |

**Supplementary Table S3: Number of SNPs showing values of $F_{ST}$, XP-EHH and ΔDAF higher than 99th percentile values of genome wide SNPs in each species.**

| Species pair | Method | Species | Number of SNPs with high value | Proportion of SNPs with high value |
|---|---|---|---|---|
| dog-cattle | fst | dog | 1199 | 0.010104 |
| | | cattle | 1107 | 0.009323 |
| dog-goat | fst | dog | 1351 | 0.010255 |
| | | goat | 1213 | 0.009274 |
| dog-horse | fst | dog | 883 | 0.010007 |
| | | horse | 876 | 0.009938 |
| dog-pig | fst | dog | 1874 | 0.009963 |
| | | pig | 1727 | 0.009281 |
| dog-sheep | fst | dog | 675 | 0.009691 |
| | | sheep | 584 | 0.008381 |
| goat-cattle | fst | goat | 3780 | 0.009531 |
| | | cattle | 3845 | 0.009604 |
| goat-pig | fst | goat | 3188 | 0.008695 |
| | | pig | 3339 | 0.009137 |
| horse-cattle | fst | horse | 1408 | 0.010219 |
| | | cattle | 1414 | 0.010252 |
| horse-goat | fst | horse | 1459 | 0.009765 |
| | | goat | 1270 | 0.008554 |
| horse-pig | fst | horse | 2047 | 0.009554 |
| | | pig | 1993 | 0.009399 |
| horse-sheep | fst | horse | 797 | 0.010128 |
| | | sheep | 723 | 0.009171 |
| pig-cattle | fst | pig | 3166 | 0.009755 |
| | | cattle | 3132 | 0.009554 |
| sheep-cattle | fst | sheep | 1181 | 0.009275 |
| | | cattle | 1305 | 0.010245 |
| sheep-goat | fst | sheep | 1432 | 0.009732 |
| | | goat | 1137 | 0.007781 |
| sheep-pig | fst | sheep | 1561 | 0.008866 |
| | | pig | 1628 | 0.00934 |
| dog-cattle | daf | dog | 1077 | 0.010526 |
| | | cattle | 1073 | 0.009785 |
| dog-goat | daf | dog | 1193 | 0.010436 |
| | | goat | 1179 | 0.009805 |
| dog-horse | daf | dog | 794 | 0.010408 |
| | | horse | 652 | 0.010193 |
| dog-pig | daf | dog | 1629 | 0.009992 |
| | | pig | 1664 | 0.00986 |
| dog-sheep | daf | dog | 622 | 0.01034 |
| | | sheep | 592 | 0.008823 |
| goat-cattle | daf | goat | 3633 | 0.010017 |
| | | cattle | 3568 | 0.009615 |
| goat-pig | daf | goat | 3131 | 0.009278 |

| | | | | |
|---|---|---|---|---|
| | | pig | 3319 | 0.009958 |
| horse-cattle | daf | horse | 1115 | 0.011031 |
| | | cattle | 1250 | 0.009886 |
| horse-goat | daf | horse | 1108 | 0.010272 |
| | | goat | 1234 | 0.009046 |
| horse-pig | daf | horse | 1499 | 0.009638 |
| | | pig | 1973 | 0.010288 |
| horse-sheep | daf | horse | 615 | 0.010729 |
| | | sheep | 666 | 0.008794 |
| pig-cattle | daf | pig | 2917 | 0.010003 |
| | | cattle | 2838 | 0.009372 |
| sheep-cattle | daf | sheep | 1135 | 0.00927 |
| | | cattle | 1185 | 0.010014 |
| sheep-goat | daf | sheep | 1427 | 0.010046 |
| | | goat | 1093 | 0.008528 |
| sheep-pig | daf | sheep | 1477 | 0.008714 |
| | | pig | 1543 | 0.00971 |
| dog-cattle | xp-ehh | dog | 1281 | 0.010836 |
| | | cattle | 1139 | 0.009597 |
| dog-goat | xp-ehh | dog | 1223 | 0.009429 |
| | | goat | 1373 | 0.010663 |
| dog-horse | xp-ehh | dog | 923 | 0.010778 |
| | | horse | 863 | 0.010078 |
| dog-pig | xp-ehh | dog | 1902 | 0.010277 |
| | | pig | 1796 | 0.009836 |
| dog-sheep | xp-ehh | dog | 685 | 0.009862 |
| | | sheep | 529 | 0.007599 |
| goat-cattle | xp-ehh | goat | 4057 | 0.010254 |
| | | cattle | 3738 | 0.009341 |
| goat-pig | xp-ehh | goat | 3680 | 0.010153 |
| | | pig | 3350 | 0.009301 |
| horse-cattle | xp-ehh | horse | 1378 | 0.010028 |
| | | cattle | 1270 | 0.009215 |
| horse-goat | xp-ehh | horse | 1374 | 0.009363 |
| | | goat | 1437 | 0.009865 |
| horse-pig | xp-ehh | horse | 1945 | 0.009292 |
| | | pig | 2023 | 0.009791 |
| horse-sheep | xp-ehh | horse | 795 | 0.010115 |
| | | sheep | 642 | 0.008149 |
| pig-cattle | xp-ehh | pig | 2980 | 0.009246 |
| | | cattle | 2931 | 0.008947 |
| sheep-cattle | xp-ehh | sheep | 1339 | 0.010526 |
| | | cattle | 1148 | 0.009017 |
| sheep-goat | xp-ehh | sheep | 1438 | 0.009781 |
| | | goat | 1359 | 0.009316 |
| sheep-pig | xp-ehh | sheep | 1512 | 0.008594 |
| | | pig | 1457 | 0.008398 |

**Supplementary Table S4: Number of SNPs showing values of $F_{ST}$, XP-EHH and ΔDAF higher than 99th percentile values of genome wide SNPs in both species.**

| pair species | methods | Number of SNPs showing high value in both species |
|---|---|---|
| dog-cattle | fst | 7 |
| dog-goat | fst | 14 |
| dog-horse | fst | 13 |
| dog-pig | fst | 24 |
| dog-sheep | fst | 8 |
| goat-cattle | fst | 40 |
| goat-pig | fst | 28 |
| horse-cattle | fst | 12 |
| horse-goat | fst | 15 |
| horse-pig | fst | 15 |
| horse-sheep | fst | 3 |
| pig-cattle | fst | 32 |
| sheep-cattle | fst | 9 |
| sheep-goat | fst | 7 |
| sheep-pig | fst | 16 |
| dog-cattle | daf | 9 |
| dog-goat | daf | 9 |
| dog-horse | daf | 6 |
| dog-pig | daf | 18 |
| dog-sheep | daf | 8 |
| goat-cattle | daf | 34 |
| goat-pig | daf | 27 |
| horse-cattle | daf | 10 |
| horse-goat | daf | 4 |
| horse-pig | daf | 9 |
| horse-sheep | daf | 4 |
| pig-cattle | daf | 23 |
| sheep-cattle | daf | 7 |
| sheep-goat | daf | 4 |
| sheep-pig | daf | 17 |
| dog-cattle | ehh | 15 |
| dog-goat | ehh | 29 |
| dog-horse | ehh | 18 |
| dog-pig | ehh | 13 |
| dog-sheep | ehh | 5 |
| goat-cattle | ehh | 45 |
| goat-pig | ehh | 34 |
| horse-cattle | ehh | 17 |
| horse-goat | ehh | 20 |
| horse-pig | ehh | 15 |
| horse-sheep | ehh | 11 |
| pig-cattle | ehh | 22 |
| sheep-cattle | ehh | 28 |
| sheep-goat | ehh | 16 |
| sheep-pig | ehh | 7 |

**Supplementary Table S5: Positively selected genes in the six Tibetan domestic mammals.**

**Supplementary Table S6: Likelihood comparison of the three demographic models for lowland population of all six species.**

| | Model | Parameters | log10(MaxL) | AIC |
|---|---|---|---|---|
| Cattle | Model A | 1 | -118437748.734 | 545425991.365 |
| | Model B | 3 | -120011411.934 | 552672982.217 |
| | Model C | 5 | -118342856.923 | 544989006.426 |
| Dog | Model A | 1 | -55920039.606 | 257521301.193 |
| | Model B | 3 | -56890197.929 | 261989049.378 |
| | Model C | 5 | -55901669.066 | 257436709.730 |
| Goat | Model A | 1 | -74825432.897 | 344583854.731 |
| | Model B | 3 | -76359253.137 | 351647361.971 |
| | Model C | 5 | -74815128.799 | 344536410.606 |
| Horse | Model A | 1 | -46507081.063 | 214173025.149 |
| | Model B | 3 | -46626246.462 | 214721806.091 |
| | Model C | 5 | -46505701.169 | 214166678.502 |
| Pig | Model A | 1 | -96387599.548 | 443881301.737 |
| | Model B | 3 | -98490162.662 | 453563966.704 |
| | Model C | 5 | -96205443.584 | 443042450.523 |
| Sheep | Model A | 1 | -138751307.110 | 638973384.770 |
| | Model B | 3 | -140518512.524 | 647111670.455 |
| | Model C | 5 | -138275998.151 | 636784514.123 |

**Supplementary Table S7. Likelihood comparison of the two joint demographic models for all six species.**

| | Model | Parameters | log10(MaxL) | AIC |
|---|---|---|---|---|
| Cattle | Model A | 11 | -170857749.390 | 786829035.536 |
| | Model B | 11 | -170874948.150 | 786908238.753 |
| Dog | Model A | 11 | -71872987.148 | 330987359.592 |
| | Model B | 11 | -71884875.910 | 331042109.364 |
| Goat | Model A | 11 | -116255337.279 | 535375635.199 |
| | Model B | 11 | -116133975.310 | 534816742.678 |
| Horse | Model A | 11 | -78891854.141 | 363310436.607 |
| | Model B | 11 | -78948549.894 | 363571530.199 |
| Pig | Model A | 11 | -144949487.218 | 667517079.011 |
| | Model B | 11 | -144941227.449 | 667479041.369 |
| Sheep | Model A | 11 | -140263779.151 | 645938595.920 |
| | Model B | 11 | -140278648.698 | 646007072.715 |

**Supplementary Table S8: Inferred parameters under the best joint demographic model of all six species**

| Parameters | Estimation | Range | | Unit |
|---|---|---|---|---|
| | | Lower bound | Upper bound | |
| Cattle | | | | |
| N ancestor of lowland | 56059 | 100 | 1.5E6 | individual |
| N bottleneck of lowland | 26763 | 100 | 2E5 | |
| N lowland | 656723 | 100 | 1.5E6 | |
| N founder of highland | 9908 | 100 | 1E5 | |
| N highland | 521945 | 100 | 1E6 | |
| T reduction of lowland | 8850 | 8750 | 11250 | year |
| T expansion of lowland | 430 | 1000 | 5000 | |
| T migration to highland | 4920 | 1000 | 5000 | |
| T expansion of highland | 335 | 100 | 5000 | |
| m lowland to highland | 4.69319e-05 | 1E-10 | 1E-1 | proportion |
| m highland to lowland | 6.05738e-07 | 1E-10 | 1E-1 | |
| Dog | | | | |
| N ancestor of lowland | 29125 | 100 | 1.5E6 | individual |
| N bottleneck of lowland | 20014 | 100 | 2E5 | |
| N lowland | 140401 | 100 | 1.5E6 | |
| N founder of highland | 6901 | 100 | 1E5 | |
| N highland | 73359 | 100 | 1E6 | |
| T reduction of lowland | 14256 | 14000 | 16000 | year |
| T expansion of lowland | 2416 | 1000 | 5000 | |
| T migration to highland | 4749 | 1000 | 5000 | |
| T expansion of highland | 1576 | 100 | 5000 | |
| M lowland to highland | 2.78471e-04 | 1E-10 | 1E-1 | proportion |
| M highland to lowland | 6.76336e-05 | 1E-10 | 1E-1 | |
| Goat | | | | |
| N ancestor of lowland | 31258 | 100 | 1.5E6 | individual |
| N bottleneck of lowland | 21987 | 100 | 2E5 | |
| N lowland | 70296 | 100 | 1.5E6 | |
| N founder of highland | 19111 | 100 | 1E5 | |
| N highland | 65996 | 100 | 1E6 | |
| T reduction of lowland | 12648 | 11000 | 13000 | year |
| T expansion of lowland | 12428 | 1000 | 13000 | |
| T migration to highland | 4908 | 1000 | 5000 | |
| T expansion of highland | 3880 | 100 | 5000 | |
| M lowland to highland | 2.87650e-07 | 1E-10 | 1E-1 | proportion |
| M highland to lowland | 5.24263e-08 | 1E-10 | 1E-1 | |
| Horse | | | | |
| N ancestor of lowland | 134713 | 100 | 1.5E6 | individual |
| N bottleneck of lowland | 205 | 100 | 2E5 | |
| N lowland | 27616 | 100 | 1.5E6 | |
| N founder of highland | 102 | 100 | 1E5 | |
| N highland | 14969 | 100 | 1E6 | |
| T reduction of lowland | 4875 | 4250 | 6750 | year |
| T expansion of lowland | 3695 | 1000 | 5000 | |
| T migration to highland | 4415 | 1000 | 5000 | |
| T expansion of highland | 2955 | 100 | 5000 | |
| M lowland to highland | 0.0021844 | 1E-10 | 1E-1 | proportion |
| M highland to lowland | 0.0052147 | 1E-10 | 1E-1 | |

| | | | | |
|---|---|---|---|---|
| Pig | | | | |
| N ancestor of lowland | 100033 | 100 | 1.5E6 | individual |
| N bottleneck of lowland | 24213 | 100 | 2E5 | |
| N lowland | 37724 | 100 | 1.5E6 | |
| N founder of highland | 11279 | 100 | 1E5 | |
| N highland | 22875 | 100 | 1E6 | |
| T reduction of lowland | 10914 | 10500 | 11500 | year |
| T expansion of lowland | 10746 | 1000 | 11500 | |
| T migration to highland | 1404 | 1000 | 5000 | |
| T expansion of highland | 1348 | 100 | 5000 | |
| M lowland to highland | 6.35171e-05 | 1E-10 | 1E-1 | proportion |
| M highland to lowland | 7.61227e-07 | 1E-10 | 1E-1 | |
| Sheep | | | | |
| N ancestor of lowland | 109261 | 100 | 1.5E6 | individual |
| N bottleneck of lowland | 38414 | 100 | 2E5 | |
| N lowland | 139041 | 100 | 1.5E6 | |
| N founder of highland | 2351 | 100 | 1E5 | |
| N highland | 68964 | 100 | 1E6 | |
| T reduction of lowland | 11244 | 10000 | 12000 | year |
| T expansion of lowland | 3716 | 1000 | 5000 | |
| T migration to highland | 4624 | 100 | 5000 | |
| T expansion of highland | 3336 | 100 | 5000 | |
| M lowland to highland | 1.66552e-04 | 1E-10 | 1E-1 | proportion |
| M highland to lowland | 4.20659e-05 | 1E-10 | 1E-1 | |

**Supplementary Table S9: Domestic time of all six species.**

| Species | Domestic date |
|---|---|
| Cattle | 8000 BCE (Wendorf and Schild 1998) |
| Dog | 13000 BCE (Thalmann et al. 2013) |
| Goat | 10000 BCE (Vigne 2011) |
| Horse | 3500 BCE (Outram et al. 2009) |
| Pig | 9000 BCE (Frantz et al. 2015) |
| Sheep | 9000 BCE (Zeder 2008) |

**Supplementary Table S10: Generation interval of all six species.**

| Species | Generation interval |
|---|---|
| Cattle | 5 years |
| Dog | 4 years |
| Goat | 4 years |
| Horse | 5 years |
| Pig | 2 years |
| sheep | 4 years |

**Supplementary Table S11: Gene enrichment analysis of differentially expressed genes after *C10orf67* knock down.** The analysis was performed by g:profiler (Reimand et al. 2011) (http://biit.cs.ut.ee/gprofiler) for enrichment analysis with Benjamini-Hochberg FDR as the multiple correction.

### Supplementary Table S12: Number of different types of SNPs at 10 different Fst bins (from 0-1). Different types of SNPs were annotated by ANNOVAR program.

| | SNP Types | 0-0.1 | 0.1-0.2 | 0.2-0.3 | 0.3-0.4 | 0.4-0.5 | 0.5-0.6 | 0.6-0.7 | 0.7-0.8 | 0.8-0.9 | 0.9-1 |
|---|---|---|---|---|---|---|---|---|---|---|---|
| Sheep | UTR | 109044 | 7266 | 2325 | 815 | 212 | 65 | 23 | 6 | 5 | 1 |
| | intronic | 11831534 | 811131 | 233672 | 77047 | 22746 | 6098 | 2230 | 619 | 197 | 82 |
| | intergenic | 29211100 | 2059407 | 581948 | 188641 | 56336 | 17076 | 4615 | 1430 | 737 | 293 |
| | splicing | 2752 | 136 | 28 | 11 | 5 | 5 | 0 | 0 | 0 | 0 |
| | upstream | 290781 | 19543 | 5974 | 2050 | 729 | 216 | 56 | 18 | 5 | 2 |
| | downstream | 276891 | 18449 | 5509 | 2017 | 693 | 168 | 44 | 9 | 2 | 1 |
| | exonic | 313402 | 19307 | 6002 | 2127 | 612 | 170 | 61 | 15 | 14 | 7 |
| | n | 133495 | 7641 | 2466 | 859 | 288 | 78 | 31 | 11 | 6 | 4 |
| | s | 177870 | 11533 | 3496 | 1259 | 320 | 91 | 28 | 4 | 8 | 3 |
| | Up/downstream | 567672 | 37992 | 11483 | 4067 | 1422 | 384 | 100 | 27 | 7 | 3 |
| dog | UTR | 103810 | 9189 | 4009 | 1220 | 519 | 173 | 40 | 14 | 1 | 0 |
| | intronic | 4949945 | 406746 | 179262 | 50860 | 22018 | 7223 | 2216 | 323 | 108 | 438 |
| | intergenic | 10471015 | 876215 | 375092 | 107157 | 47543 | 15042 | 3822 | 855 | 358 | 387 |
| | splicing | 861 | 50 | 27 | 5 | 2 | 4 | 0 | 0 | 0 | 0 |
| | upstream | 122726 | 11258 | 4979 | 1587 | 726 | 242 | 89 | 18 | 11 | 17 |
| | downstream | 117498 | 10143 | 4529 | 1345 | 546 | 193 | 70 | 33 | 12 | 11 |
| | exonic | 111678 | 9604 | 4354 | 1334 | 578 | 232 | 63 | 20 | 11 | 12 |
| | n | 49073 | 3939 | 1735 | 520 | 238 | 101 | 25 | 7 | 8 | 6 |
| | s | 61948 | 5626 | 2599 | 803 | 338 | 128 | 38 | 13 | 3 | 6 |
| | Up/downstream | 240224 | 21401 | 9508 | 2932 | 1272 | 435 | 159 | 51 | 23 | 28 |
| cattle | UTR | 94304 | 10249 | 3790 | 1373 | 412 | 173 | 59 | 11 | 0 | 0 |
| | intronic | 9976200 | 1055732 | 380112 | 141658 | 53056 | 18099 | 5672 | 1363 | 150 | 12 |
| | intergenic | 26422240 | 2900024 | 1060427 | 408790 | 154565 | 52303 | 15896 | 3732 | 552 | 99 |
| | splicing | 1419 | 137 | 53 | 16 | 9 | 1 | 0 | 2 | 0 | 0 |
| | upstream | 256525 | 27693 | 10337 | 3632 | 1195 | 418 | 147 | 40 | 5 | 0 |
| | downstream | 262599 | 27410 | 10171 | 3745 | 1299 | 493 | 127 | 34 | 5 | 0 |
| | exonic | 262391 | 27636 | 10294 | 3613 | 1228 | 440 | 131 | 56 | 6 | 1 |
| | n | 102543 | 10817 | 4041 | 1382 | 489 | 169 | 50 | 26 | 2 | 1 |
| | s | 158637 | 16686 | 6218 | 2219 | 737 | 266 | 80 | 30 | 4 | 0 |
| | Up/downstream | 519124 | 55103 | 20508 | 7377 | 2494 | 911 | 274 | 74 | 10 | 0 |
| horse | UTR | 15610 | 986 | 229 | 58 | 13 | 1 | 1 | 0 | 0 | 0 |
| | intronic | 4449137 | 279655 | 64960 | 12866 | 2274 | 363 | 33 | 2 | 3 | 0 |
| | intergenic | 12055843 | 744408 | 168933 | 33187 | 5880 | 1020 | 166 | 16 | 2 | 0 |
| | splicing | 1168 | 45 | 14 | 2 | 0 | 0 | 0 | 0 | 0 | 0 |
| | upstream | 148631 | 9298 | 2287 | 498 | 114 | 21 | 5 | 0 | 0 | 0 |
| | downstream | 140802 | 8862 | 2098 | 401 | 74 | 15 | 2 | 1 | 0 | 0 |
| | exonic | 128963 | 7913 | 1905 | 385 | 88 | 20 | 2 | 0 | 0 | 0 |
| | n | 59301 | 3480 | 860 | 159 | 38 | 11 | 1 | 0 | 0 | 0 |
| | s | 68945 | 4390 | 1036 | 224 | 50 | 8 | 1 | 0 | 0 | 0 |
| | up/downstream | 289433 | 18160 | 4385 | 899 | 188 | 36 | 7 | 1 | 0 | 0 |
| pig | UTR | 219947 | 8600 | 1452 | 301 | 80 | 25 | 5 | 2 | 0 | 0 |
| | intronic | 11642167 | 435452 | 69627 | 11945 | 2659 | 525 | 112 | 36 | 7 | 2 |
| | intergenic | 32734841 | 1184921 | 182165 | 30433 | 8375 | 1600 | 389 | 90 | 23 | 4 |
| | splicing | 1478 | 51 | 9 | 4 | 0 | 0 | 0 | 0 | 0 | 0 |

| | | | | | | | | | | | |
|---|---|---|---|---|---|---|---|---|---|---|---|
| | upstream | 329878 | 13669 | 2396 | 593 | 142 | 39 | 6 | 2 | 0 | 0 |
| | downstream | 341164 | 13283 | 2278 | 396 | 119 | 24 | 3 | 3 | 0 | 1 |
| | exonic | 270278 | 11836 | 2056 | 432 | 130 | 26 | 8 | 2 | 0 | 0 |
| | n | 96624 | 3836 | 630 | 121 | 36 | 4 | 1 | 1 | 0 | 0 |
| | s | 171739 | 7935 | 1417 | 308 | 93 | 22 | 7 | 1 | 0 | 0 |
| | up/downstream | 671042 | 26952 | 4674 | 989 | 261 | 63 | 9 | 5 | 0 | 1 |
| goat | UTR | 311433 | 8202 | 1431 | 345 | 78 | 23 | 1 | 0 | 0 | 0 |
| | intronic | 19161175 | 541651 | 111554 | 25814 | 5530 | 1078 | 380 | 84 | 28 | 11 |
| | intergenic | 34992134 | 1087557 | 221311 | 48375 | 9800 | 2058 | 567 | 184 | 84 | 23 |
| | splicing | 4303 | 114 | 15 | 8 | 1 | 1 | 0 | 0 | 0 | 0 |
| | upstream | 320435 | 9270 | 1926 | 410 | 97 | 20 | 4 | 0 | 1 | 0 |
| | downstream | 346977 | 9644 | 1950 | 464 | 113 | 23 | 5 | 1 | 0 | 0 |
| | exonic | 626088 | 16909 | 2551 | 470 | 131 | 23 | 3 | 2 | 1 | 0 |
| | n | 251466 | 6729 | 1166 | 233 | 58 | 10 | 1 | 1 | 0 | 0 |
| | s | 348202 | 9490 | 1289 | 214 | 63 | 12 | 1 | 1 | 1 | 0 |
| | up/downstream | 667412 | 18914 | 3876 | 874 | 210 | 43 | 9 | 1 | 1 | 0 |

**Supplementary Table S13:** Examples of positively selected genes harboring non-synonymous SNPs display high level of population differentiation between Tibetan domestic mammals and lowland animals ($F_{ST}$>0.5).

| Species | Genes | Function | Sites |
|---|---|---|---|
| cattle | *PDCD7* | apoptosis | P16A |
| cattle | *DIDO1* | apoptosis | D895E, H1627R, R2094H |
| cattle | *GAK* | apoptosis | V503A |
| cattle | *CARD14* | apoptosis | F497S |
| cattle | *EPAS1* | hypoxia adaptation | P362L |
| cattle | *HMGA2* | height | A64P |
| cattle | *APOB* | lipid metabolism | L1436H, N2212H |
| cattle | *ALDH1L2* | metabolism | I74L |
| cattle | *IDO2* | metabolism | R402L |
| cattle | *ADH7* | metabolism/height | L42P, S52P, C58R |
| cattle | *ERMARD* | nervous system | V579A, R647H |
| cattle | *ADCY7* | nervous system | R713W |
| dog | *EPAS1* | hypoxia adaptation | G305S, D494E, V500M, P750S |
| dog | *NLRP1* | apoptosis | R49H, A1347T, R789T |
| dog | *CXCL16* | apoptosis | R180P |
| dog | *XAF1* | apoptosis | Y35H |
| dog | *ALOX15* | lipid metabolism | R349C |
| dog | *MSRB3* | energy metabolism | T166A, G179S |
| dog | *PER1* | nervous system | p.R966C, |
| dog | *SUFU* | nervous system | p.V228I, |
| dog | *CDH8* | nervous system | p.T414M, |
| horse | *ATG2B* | lipid metabolism | R374G |
| pig | *AATK* | apoptosis | L805V |
| pig | *SLC25A47* | energy metabolism | A144T |
| sheep | *RPAP3* | apoptosis | C3R |
| sheep | *CASP8AP2* | apoptosis | K1738R |
| sheep | *GRAMD4* | apoptosis | p.T415M, |
| sheep | *NDUFB1* | energy metabolism | G17V, W39R |
| sheep | *MDN1* | energy metabolism | L545F, G2991D, S2994A, T2967I, N2879D |
| sheep | *CA4* | energy metabolism | N190T |
